# RS-FISH: Precise, interactive, fast, and scalable FISH spot detection

**DOI:** 10.1101/2021.03.09.434205

**Authors:** Ella Bahry, Laura Breimann, Marwan Zouinkhi, Leo Epstein, Klim Kolyvanov, Xi Long, Kyle I S Harrington, Timothée Lionnet, Stephan Preibisch

**Author notes:** equal contribution (Ella Bahry, Laura Breimann, Marwan Zouinkhi).

## Abstract

Fluorescent in-situ hybridization (FISH)-based methods are powerful tools to study molecular processes with subcellular resolution, relying on accurate identification and localization of diffraction-limited spots in microscopy images. We developed the Radial Symmetry-FISH (RS-FISH) software that accurately, robustly, and quickly detects single-molecule spots in two and three dimensions, making it applicable to several key assays, including single-molecule FISH (smFISH), spatial transcriptomics, and spatial genomics. RS-FISH allows interactive parameter tuning and scales to large sets of images as well as tera-byte sized image volumes such as entire brain scans using straight-forward distributed processing on workstations, clusters, and in the cloud.

## Main

New FISH-based imaging methods are continuously developed to gain insights into cellular processes, for example, by resolving the subcellular localization of single RNA molecules ^1,2^ or subnuclear 3D arrangement of DNA regions ^3,4^. Classically, smFISH was used to visualize individual mRNA molecules for single genes in small samples ^1,2^. New methods that employ probe amplification, probe multiplexing, or barcodes drive the fields of spatial transcriptomics and spatial genomics, enabling the subcellular visualization of thousands of genes with single-molecule sensitivity in complex tissues ^5–10^, as well as entire chromosomes with high-resolution at nanometer scale ^3^.

Extracting information from smFISH, spatial transcriptomics, or spatial genomics images relies on the precise detection of diffraction-limited spots. Important properties of spot detection software are accuracy and speed of detection, as well as being accessible to experimentalists. Recently, scalability to large datasets is becoming important since the detection of subtle transcriptional changes relies on the analysis of thousands of smFISH images ^11,12^, increasingly large samples in the tera-byte range are being imaged ^13^, and spatial transcriptomics methods are applied to increasingly large samples with many rounds of sequential hybridization and imaging (**Fig. 1, Supplement**). Several methods are available to the community, however, all commonly used packages do not allow interactive parameter tuning, which makes their application tedious for experimentalists, and do not scale to large datasets due to missing options for straight-forward automation and distribution, and partly because of slower processing times ^1,14–17^. To overcome these restrictions, we developed RS-FISH that uses an extension of Radial Symmetry^18^ (RS) to robustly and quickly identify single-molecule spots in 3D with high precision (**Fig. 1a**). RS-FISH can be run as an interactive, scriptable Fiji plugin^19^, as a command-line tool, and as a cluster and a cloud distributable package for large volumes or for datasets consisting of thousands of images (**Fig. 1g, h**).

**Figure 1:**
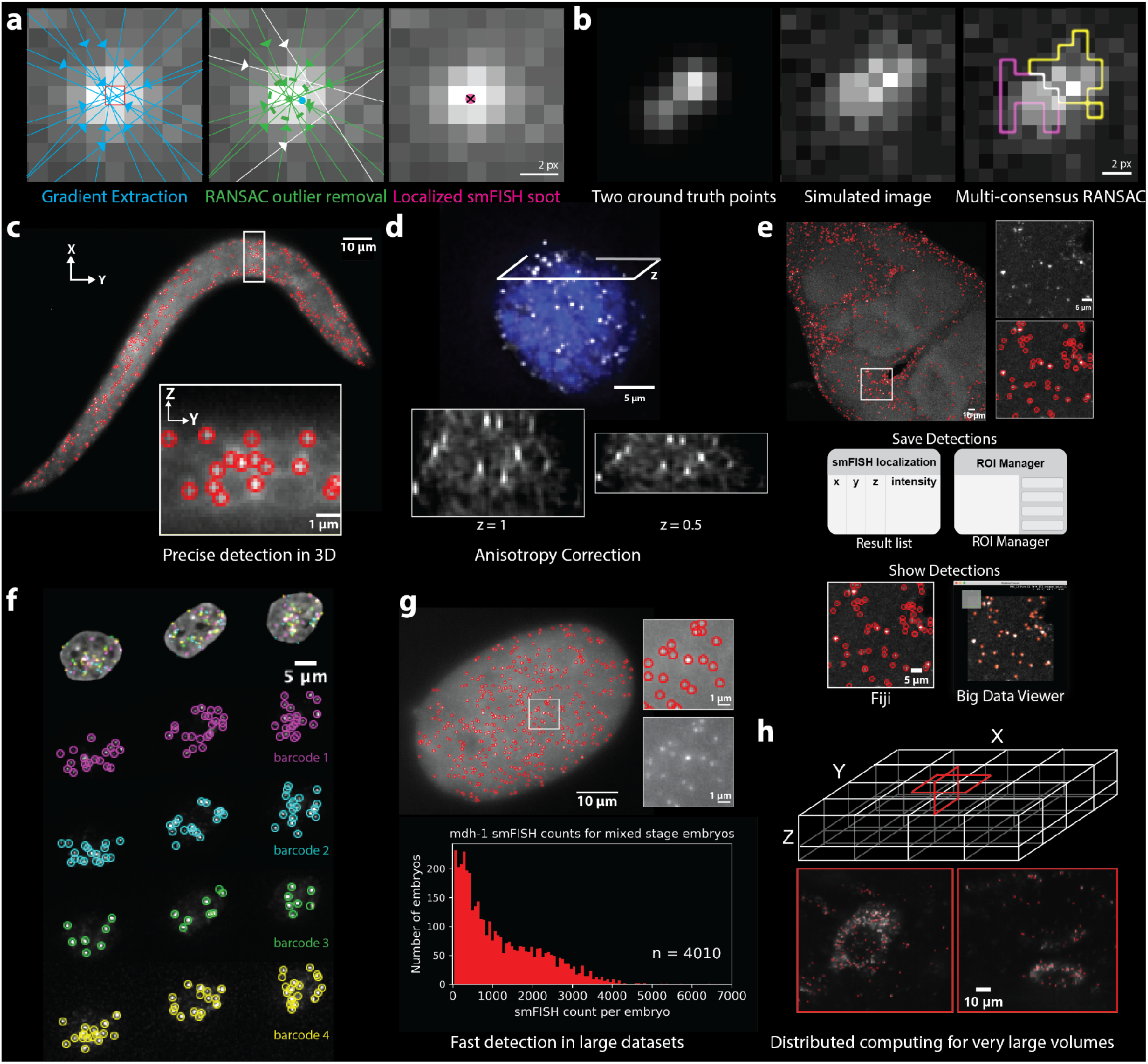
RS-FISH accurately detects smFISH spots. **(a)** Illustration of the concept of single smFISH spot detection using Radial Symmetry-based RANSAC. (left) Blue lines indicate the gradients calculated in a local patch around a DoG-detected location (red square) for radial symmetry fitting. Arrowheads indicate the location for which example gradients are shown. (middle) Gradients that agree on a common center point (green gradients, green dot) given a defined error (green dotted line) are identified using RANSAC outlier removal. Using all gradients would lead to a different center point (blue). (right) RS-FISH detects the center of the smFISH spot by computing the intersection point of all inlier gradients (pink dot with the black cross). **(b)** Detection of two close points using multi-consensus RANSAC. (left) Two ground truth points in close proximity are shown without noise. (middle) Both points are detected as a single spot during DoG due to simulated low SNR. (right). Using the multi-consensus RANSAC, the area was identified to consist of two independent detections. The two sets of gradients (i.e., pixels) used to detect each spot are highlighted in yellow and pink. **(c)** Detection of spots is possible on 2D or 3D images. XY image of a single z-slice of a *C. elegans* larva expressing *lea-1* labeled with smFISH detected. Red circles highlight the RS-FISH detected spots. ZY image of the highlighted area in the worm. **(d)** To optimize the detection of radial centers, the anisotropy of the image can be determined and corrected using a global scale factor. The example image shows a mouse embryonic stem cell labeled by smFISH for cdx-2 mRNA. **(e)** Possibilities for downstream analysis using RS-FISH. The detections can be saved as a Results table or directly transferred to the ROI manager. The detected spots in the Results table can be shown in Fiji or using the BigDataViewer for visual inspection. Example images show a max projection of 5 z-slices of a Drosophila brain with smFISH labeled for Pura mRNA^27^. **(f)** RS-FISH can detect other diffraction-limited spots. PGP1f cells labeled with OligoFISSEQ using barcodes with four different fluorophores^3^. Images were published previously by Nguyen et al.^3^ and were kindly provided to us. Single spots detected by RS-FISH were labeled in 4 different colors. **(g)** RS-FISH allows fast detection in large datasets through batch processing. Image of an early *C. elegans* embryo with smFISH labeling for *dpy-23* mRNAs. Distribution of mRNA counts for 4010 mixed stage single embryos for *mdh-1* mRNA. **(h)** The spark version of RS-FISH can analyze large N5 volumes in a distributed fashion locally, on the cluster, or in the cloud. Example images of Expansion-Assisted Iterative Fluorescence In Situ Hybridization (EASI-FISH) spots detected in a 148 GB lightsheet image of a tissue section of the lateral hypothalamic area for the gene Map1b^13^. The data was previously published by Wang et al.^13^ and was kindly provided to us.

RS is an efficient, non-iterative alternative to accurate point localization using Gaussian fitting that was developed for localizing two-dimensional circular objects by computing the intersection point of image gradients (**Fig. 1a**)^18^. We first derived a 3D version of the RS method similar to Liu et al. ^20^ (**Methods**) that additionally extends to higher dimensions, which has potential for spatiotemporal localization of blinking 3D spots. Second, we extended RS to support axis-aligned, ellipsoid objects without the need for scaling the image^20^, enabling RS-FISH to account for typical anisotropy in 3D microscopy datasets that results from different pixel sizes and point spread functions in the lateral (x,y) compared to the axial (z) dimensions (**Fig. 1d, Methods**). Third, the computation speed of RS allowed us to combine RS with robust outlier removal using Random Sample Consensus^21^ (RS-RANSAC) to identify sets of image gradients that support the same ellipsoid object given a specific error for the gradient intersection point (**Methods**). This allows RS-FISH to identify sets of pixels that support a user-defined localization error for individual spots (**Fig. 1a**), separate close detections (**Fig. 1b**), and ignore outlier pixels that disturb localization (e.g., dead or hot camera pixels).

RS-FISH first generates a set of seed points by thresholding the Difference-of-Gaussian (DoG)^22^ filtered image to identify potential locations of diffraction-limited spots. Next, image gradients are extracted from local pixel patches around each spot, which are optionally corrected for non-uniform fluorescence backgrounds. Before RS localization, gradients are rescaled along the axial dimension to correct for dataset anisotropy using an anisotropy factor that depends on pixel spacing, resolution, and point spread function. The anisotropy factor can be computed from the microscopy image itself and does not change while acquisition parameters are held constant (**Fig. 1d, Methods**). Optionally, RS-RANSAC can be run in multi-consensus mode, which allows detecting spots that were too close for the DoG detector to separate them during seed point generation (**Fig. 1b, Methods**). Finally, to avoid potentially redundant detections, detections are filtered to be at least 0.5 pixels apart from each other. Each spot’s associated intensity values are computed using either simple linear interpolation at the spot’s sub-pixel location or by fitting a Gaussian to the subset of pixels that support the spot as identified by RS-RANSAC.

RS-FISH pixel operations are implemented in ImgLib2^23^, and RS fitting and RS-RANSAC are implemented using the image transformation framework mpicbg^24^. All operations can be executed in blocks allowing straight-forward parallelization where compute effort scales linearly with the size of the data up to the petabyte range (**Methods**). Importantly, RS-FISH’s parameters can be interactively tuned on small and large datasets using the Fiji plugin. Once the right set of parameters is identified, RS-FISH can be run and macro-scripted in Fiji or can be executed in a scriptable mode for straight-forward parallel execution on compute clusters or cloud services (e.g., Amazon Web Services (AWS)) using Apache Spark, for which we provide example scripts including resaving into the N5 file format (**Fig. 1g, Supplement**). The results are saved as CSV or can be transferred to the ROI manager for downstream analysis in Fiji (**Fig. 1e, Supplement**). The saved point clouds can be overlaid onto the images using Fiji^19^ or BigDataViewer^25^ for interactive visual inspection of even very large datasets (**Fig. 1h, Supplementary Video 1**).

To validate and benchmark RS-FISH, we performed quantitative comparisons against FISH-quant^14^, Big-FISH^26^, AIRLOCALIZE^17^, Starfish^16^, and deepBlink^15^ using (i) simulated smFISH images with varying noise levels to assess detection performance, (ii) real *C. elegans* embryo datasets for runtime measurements, and (iii) large lightsheet datasets^13^. We show that RS-FISH is on par with the best methods in terms of detection performance (**Fig. 2a-c, Supplement**) while running 3.8x – 7.1x times faster than established methods (**Fig. 2d, Supplement**). We highlighted the ability to easily parallelize RS-FISH on the cloud by running smFISH extraction on 4.010 *C. elegans* image stacks (in total ∼100 GB) in 18 minutes on AWS at the cost of 18,35 USD (**Fig. 1g**). Importantly, RS-FISH is currently the only method that can be directly applied to large volumes (**Fig. 1h**). Processing a reconstructed 148 GB lightsheet image stack took 32 CPU hours (∼1 hour on a modern workstation). In comparison, a complex wrapping software for distributing AIRLOCALIZE, specifically developed for the EASI-FISH project to run on the HHMI Janelia cluster, required significant development effort and took 156 CPU hours to finish the same task. We developed RS-FISH based on a generic derivation of 3D Radial Symmetry for anisotropic objects that is efficiently implemented using ImgLib2, Fiji, and Spark. RS-FISH runs as a Fiji plugin allowing interactive parameter adjustment and result verification on small and large images, making the task of correctly detecting most diffraction-limited spots in microscopy images as accessible as possible to experimentalists. Processing speed is significantly improved while achieving similar localization performance to established methods. RS-FISH is simple to install and run through Fiji, additionally providing macro-recording functionality to automate FISH spot detection easily. Our efficient block-based implementation allows easy single-molecule spot detection in large datasets or big volumes using local processing, clusters, or the cloud. Importantly, while we only show a 148 GB dataset, there is no conceptual limit that prohibits RS-FISH from being executed on significantly larger volumes well into the petabyte range. Taken together, RS-FISH is an accurate, easy-to-use, versatile, and scalable tool that makes FISH spot detection on small and especially large datasets amenable to experimentalists and whose functionality extends to the dynamically growing fields of spatial transcriptomics and spatial genomics.

**Figure 2:**
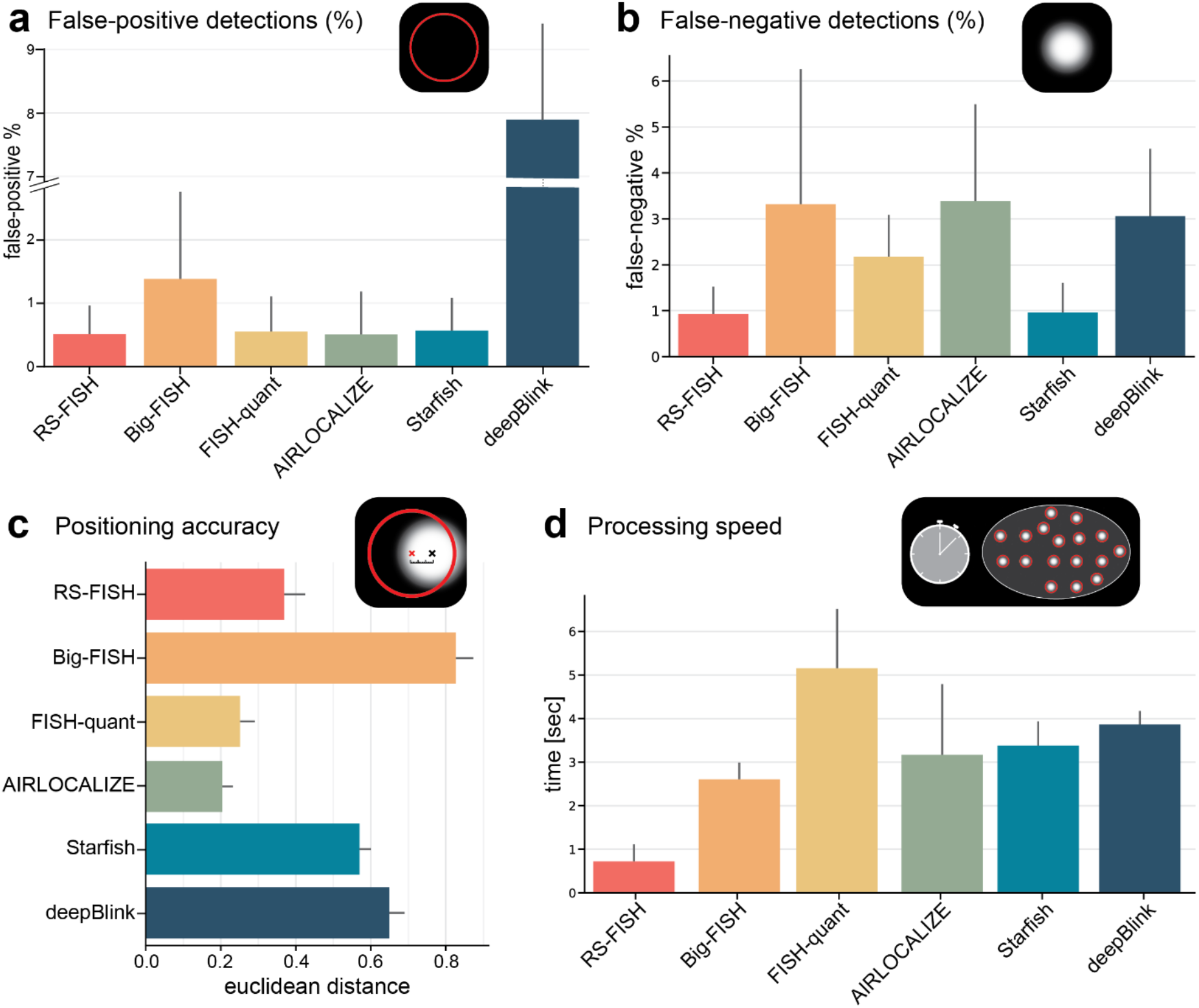
Performance of different spot localization tools. **(a)** False-positive detection percentage for wrongly detected spots using simulated data. **(b)** False-negative detection percentage for missed detections using simulated data. **(c)** Localization accuracy is measured as the Euclidean distance (pixels) between the detected spot and the true center of simulated spots. **(d)** Median processing speed for thirteen 3D smFISH images of *C. elegans* embryos. Details on running parameters can be found in the Supplementary Material.

## Methods

### *n*-dimensional derivation of Radial Symmetry localization

The goal of RS is to accurately localize a bright, circular spot ***p***_***c***_ with subpixel accuracy. In noise-free data, image gradients **∇*I*(*p***_***k***_)at locations ***p***_***k***_ point towards the center of the spot and intersect in that single point ***p***_***c***_ (**Fig. 1a**), thus computing the intersection point solves the problem of accurate localization. In realistic images that contain noise, these gradients do not intersect, therefore computing ***p***_***c***_ constitutes an optimization problem that RS solved using least-squares minimization of the distances ***d***_***k***_ between the common intersection point ***p***_***c***_ and all gradients **∇ *I* (*p***_***k***_ ) (**Supplementary Figure 1**).

We extend RS to 3D similar to Liu et al. ^20^ and additionally describe how to generalize the derivation to the n-dimensional case. To achieve this, we replace the Roberts cross operator with separable convolution for image gradient **∇ *I* (*p***_***k***_) computation, and we use vector algebra to compute the intersection point ***p***_***c***_ of image gradients. The derivations are shown in detail in **Supplementary Figure 1** and **Supplemental Material**.

### Radial Symmetry for axis-aligned ellipsoid (non-radial) objects

Diffraction-limited spots in 3D microscopy images are usually not round but show a scaling in the axial (z) dimension compared to the lateral (xy) dimensions. Previous solutions suggested scaling the image in order to be able to detect spots using RS ^20^. This can be impractical for large datasets, and it might affect localization quality as the image intensities need to be interpolated for scaling. Here, we extend the RS derivation to directly compute the intersection point ***p***_***c***_ from anisotropic images by applying a scale vector ***s*** to point locations ***p***_***k***_ and applying the inverse scale vector ***s***^***-1***^ to the image gradients **∇ *I* (*p***_***k***_). While we derive the case specifically for 3D, it can be straight-forward applied to higher dimensions. The derivation is shown in detail in the **Supplemental Material**.

RS-FISH supports a global scale factor (called anisotropy factor) for the entire dataset that compensates for anisotropy of the axial (z) dimension, which can be computed from an image containing diffraction-limited spots (**Supplement**).

### Radial Symmetry RANSAC

RS localization is implemented as a fast, closed-form solution, and it is therefore feasible to combine it with robust outlier removal. We use Random Sample Consensus (RANSAC)^21^ in order to identify the maximal number of gradients **∇ *I* (*p***_***k***_) that support the same center point ***p***_***c***_ given a maximal distance error ***ε*** so that all ***d***_***k***_ < ***ε***.

To achieve this, RANSAC randomly chooses the minimal number of gradients (i.e., two gradients) from the set of all gradients (candidate gradients) to compute the center point and tests how many other gradients fall within the defined error threshold ***ε***. This process is repeated until the maximal set of gradients is identified (inlier gradients) and the final center point ***p***_***c***_ is computed using all inlier gradients. This allows RS-FISH to exclude artifact pixels and to differentiate close-by spots.

To identify and locate close-by points, RS-FISH runs a multi-consensus RANSAC. Here, RANSAC is run multiple times on the same set of candidate gradients. After each successful run that identifies a set of inliers, these inliers are removed from the set of candidate gradients, and RANSAC tries to identify another set of inliers (**Fig. 1b**). This process is iterated until no other set of inliers (corresponding to a FISH spot) can be found in the local neighborhood of each DoG spot. To not detect random noise, the minimal number of inliers required for a spot can be adjusted (typically around 30).

### Implementation details and limits

RS-FISH is implemented in Java using ImgLib2, the *mpicbg* framework, BigDataViewer, Fiji, and Apache Spark. The computation of RS is performed in blocks with a size of *p*_*d*_ for each dimension *d* (e.g., 256×256×128 pixels) and requires an overlap of only one pixel in each dimension with neighboring blocks, thus the overhead 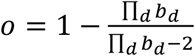 is minimal (e.g. 1.5% for 256×256×256 blocks or 0.6% for 1024×1024×1024 blocks). When processing each block the local process has access to the entire input image which is either held in memory when running within Fiji or is lazy-loaded from blocked N5 datasets when running on large volumes using Apache Spark. Since the computation across blocks is embarrassingly parallel, computation time linearly scales with the dataset size. Thus, RS-FISH will run on very large volumes supported by N5 and ImgLib2. Due to current limitations in Java arrays the theoretical upper limit is 2^31^ = 2.147.483.648 blocks, each block maximally containing 2^31^ = 2.147.483.648 pixels (e.g. 2048×2048×512 pixels). Given sufficient storage and compute resources, the limit for RS-FISH is thus 4072 peta-pixels (4072 petabyte @ 8bit or 8144 petabyte @ 16bit) taking into account the overhead, whereas every individual block locally only processes 2 gigapixels (2^31^ = 2.147.483.648 pixels).

The code can be executed on an entire image as a single block for smaller images, or in many blocks multi-threaded or distributed using Apache Spark. It is important to note that RS-RANSAC uses random numbers to determine the final localization of each spot. We use fixed seeds for initializing each block, therefore the results for a single block of the same size in the same image with the same parameters are constant. However, if one compares results of blocks of different sizes (e.g. single-threaded vs. multi-threaded), the results will be slightly different as the RANSAC-based localizations are not traversing the DoG maxima in the same order, thus initializing RANSAC differently. For practicality, the interactive Fiji mode only runs in single-threaded mode (although the DoG image is computed multi-threaded) to yield comparable results across different testing trials. Importantly, this only applies if RANSAC mode is used for localization. Multi-threaded processing is available in the recordable advanced mode in the Fiji plugin, while the Apache Spark based distribution can be called from the corresponding RS-FISH-Spark repository.

### Data Simulation for assessing localization performance

In order to create ground-truth datasets for assessing localization performance, we generated images simulating diffraction-limited spots in the following way: (*x,y,z*) spot positions were randomly assigned within the z-stack chosen dimensions, and each spot was assigned a brightness picked from a normal distribution. We computed the intensity *I* (*x,y,z*)generated by each spot as follows: we first computed the predicted average number of photons received by each pixel *I*_*pred*_(*x,y,z*) computed using a gaussian distribution centered on the spot, with user-defined lateral and axial extensions. We then simulated the actual intensity collected at each pixel using a Poisson-distributed value with mean *I*_*pred*_ (*x,y,z*). We eventually added gaussian-distributed noise to each pixel of the image.

Code for generating the images with simulated diffraction-limited spots is available in the GitHub repository. There is also a folder included with the simulated data used in the parameter grid search and benchmarking. https://github.com/PreibischLab/RS-FISH/tree/master/documents/Simulation_of_data

### Benchmarking RS-FISH against commonly used spot detection tools

RS-FISH performance was benchmarked against the leading tools for single-molecule spot detection in images. The tools compared in the benchmarking are FISH-quant^14^ (Matlab), Big-FISH ^26^ (Python), AIRLOCALIZE ^17^ (Matlab), Starfish^16^ (Python), and deepBlink^15^ (Python, TensorFlow). Localization performance comparison was done on simulated images with known ground truth spot locations, and computation time comparison was performed using real three-dimensional *C. elegans* smFISH images. We created a dedicated analysis pipeline for each tool to test localization performance and compute time. For localization performance comparison, a grid search over each tool’s pipeline parameter space was run (excluding deepBlink, as a pre-trained artificial neural network was used, more details regarding deepBlink are discussed in **Supplemental Notes**). Importantly, tools use different offsets for their pixel coordinates, which depends on the respective pixel origin convention (e.g. does a pixel lie at 0,0 or 0.5,0.5, does the z index start with 0 or 1). In our benchmarks we detect these offsets by computing the precision (the average, signed per-dimension difference between predicted and ground-truth spots) and corrected for these offsets if necessary. Notably, the Matlab tools AIRLOCALIZE and FISH-quant both show a 0.5 pixel offset in *xy*, and 1 pixel in *z* compared to the other tools. For computation time comparison, each pipeline’s parameters were selected to produce a similar number of detected spots for each image.

The comparison shows that RS-FISH is on par with currently available spot detection tools in localization performance (**Fig. 2a-c, Supplement**) while running 3.8x – 7.1x times faster (**Fig. 2d, Supplement**). Additionally, RS-FISH is currently the only available tool that can be directly applied to large images, which we highlight using a 148 GB lightsheet image stack^13^ (**Fig 1h, Supplementary Video 1, Supplement)**. The image size of the lightsheet stack is 7190×7144×1550 pixels, and the block size used for detection was 256×256×128 pixels. The detection of spots using RS-FISH took 3263 sec (∼32 CPU hours) for the entire image on a 36 CPU workstation with 2x Intel Xeon Gold 5220 Processor@2.2Ghz. The runtime cannot be directly compared to the custom extension of AIRLOCALIZE that was developed for the same project as it is written to specifically run only on the Janelia cluster. The compute time of 156 CPU hours was extracted from the cluster logs of the submission scripts and was executed on a mix of Intel SkyLake (Platinum 8168) @2.7GHz and Intel Cascade Lake (Gold 6248R) @3.0Ghz CPUs. The overall speed increase of ∼5x generally agrees with our measurements in **Fig. 2d** and the performance of a mix of these CPUs is comparable to the workstation CPUs (according to https://www.cpubenchmark.net). Importantly, RS-FISH runs on such volumes natively and can easily be executed on a cluster or in the cloud, thus easily scales to significantly larger datasets. At the same time, the AIRLOCALIZE implementation is limited to the Janelia cluster, but could be extended to other LSF clusters that support job submission.

Benchmarking analysis details are in the **Supplemental Material**, and all scripts and complete documentation are in the RS-FISH GitHub repository.

## Data availability

All datasets used for benchmarking are available in the RS-FISH GitHub repository, which includes simulations and 3D smFISH images of *C. elegans* embryos.

## Code availability

RS-FISH is implemented as open-source in Java/ImgLib2 and provided as a macro-scriptable Fiji plugin and stand-alone command-line application capable of cluster and cloud execution. Code source, tutorial, documentation, and example images are available at: https://github.com/PreibischLab/RS-FISH and https://github.com/PreibischLab/RS-FISH-Spark.

## Acknowledgments

We would like to thank Stephan Saalfeld for providing the *mpicbg* framework, the System Biology imaging platform, and Andrew Woehler for technical support. Dhana Friedrich for discussions about the *C. elegans* smFISH imaging protocol, Florian Mueller and Arthur Imbert for their advice on running Big-FISH, and Jeffrey Chao and Bastian Eichenberger on their referral to deepBlink’s 3D solution. Additional datasets for images in the main figure were kindly provided by Huy Quoc Nguyen, Shyamtanu Chattoraj and Ting Wu, and Yuhan Wang from the Multifish team project. We would like to thank Rob Klose; the mESC smFISH were labeled and imaged in his lab by L.B.

E.B. was supported by the HFSP grant RGP0021/2018-102 and MDC-Berlin, L.B. was supported by MDC-Berlin and the Joachim Herz Foundation (#850022), L.E. and K.H were supported by MDC-Berlin and the Helmholtz Imaging Platform, K.K. was supported by MDC-Berlin, T.L. was supported by NIH grant R01 GM127538 and Melanoma Research Foundation Team Science award # 687306, and S.P. was funded by MDC-Berlin and HHMI Janelia.

