## Supplementary Note for "RS-FISH: Precise, interactive, fast, and scalable FISH spot detection"

Supplementary Figure:

1. Concept of 3D Radial Symmetry

Supplementary Tables:

1. Comparison of localization performance on simulated data
2. Comparison of execution time of real 3D datasets
3. Information on datasets used for this study

Supplementary Videos:

1. Fly-through video of a large lightsheet dataset with RS-FISH detected spots using BigDataViewer.

Supplementary Notes:

1. Derivation of the three-dimensional Radial Symmetry localization
2. Derivation for supporting axis-aligned ellipsoid (non-radial) objects
3. Computing the global scale factor
4. Accuracy benchmarking RS-FISH against commonly used tools
5. Computation time benchmarking
6. Tutorial RS-FISH
7. Batch processing and macro tutorial
8. Distributed processing using RS-FISH-Spark
9. smFISH protocol for *C. elegans*
10. smFISH protocol for mouse ES cells using cytospin
11. smFISH protocol for cleared whole-mount adult Drosophila brains

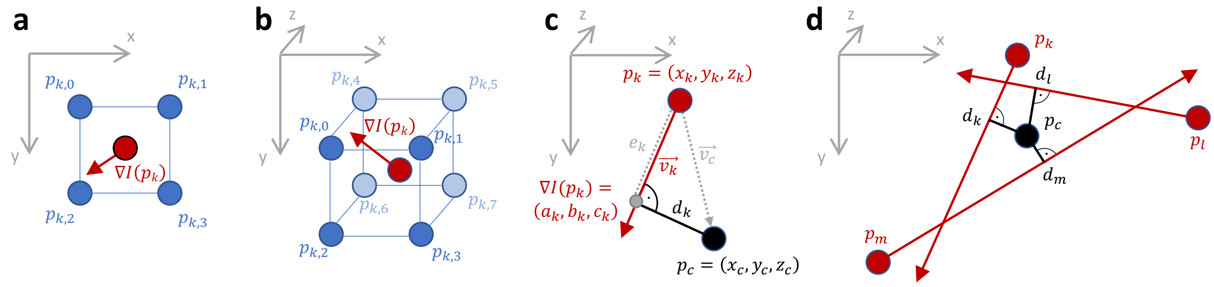


**Supplementary Figure 1**: *Concept of 3D Radial Symmetry, derived from Parthasarathy [1]. (**a**) illustrates for the two-dimensional case the concept of the intensity gradient $\nabla I(p)$ at the half-pixel offset locations $p$, computed from four intensity values $I(p)$ at locations $\{p_0, p_1, p_2, p_3\}$ that intersects with the intensity maximum. (**b**) generalizes the concept to three dimensions, which requires eight intensity values $I(p)$ at locations $\{p_0, \ldots, p_7\}$. (**c**) Parthasarathy defined the center point $p_c = (x_c, y_c, z_c)$ and the shortest distance to $\nabla I(p)$ as $d_k$, which directly extends to 3D. (d) Illustrates that minimizing $\sum_i d_i$ yields the common intersection point $p_c$.*

**Supplementary Table 1**: Comparison of localization performance on simulated data. The metrics used are the mean Euclidean distance (pixels) between the ground truth location to each detected spot, the percentage of missed detections (FN), and the percentage of false detections (FP). The tools compared are AIRLOCALIZE (Air), Big-FISH (BF), deepBlink (DB), FISH-Quant (FQ), RS-FISH (RS), and Starfish (SF).

| **Metric** | **Euclidean Distance** | | | | | | **FN Percent** | | | | | | **FP Percent** | | | | | |
|---|---|---|---|---|---|---|---|---|---|---|---|---|---|---|---|---|---|---|
| **Method** | **Air** | **BF** | **DB** | **FQ** | **RS** | **SF** | **Air** | **BF** | **DB** | **FQ** | **RS** | **SF** | **Air** | **BF** | **DB** | **FQ** | **RS** | **SF** |
| **Image** | | | | | | | | | | | | | | | | | | |
| Empty Bg Density Range Sigxy 1 SigZ 2/Poiss_300spots_bg_200_2_I_300_0_img0 | 0.2 | 0.7 | 0.5 | 0.2 | 0.3 | 0.6 | 2.3 | 1.3 | 4.7 | 2.3 | 1 | 1 | 0.3 | 0 | 4.3 | 0 | 0 | 0 |
| Empty Bg Density Range Sigxy 1 SigZ 2/Poiss_300spots_bg_200_2_I_300_0_img1 | 0.2 | 0.6 | 0.5 | 0.2 | 0.3 | 0.5 | 2 | 0.3 | 4 | 1.3 | 0 | 0.3 | 0.3 | 0 | 2.3 | 0 | 0 | 0 |
| Empty Bg Density Range Sigxy 1 SigZ 2/Poiss_300spots_bg_200_2_I_300_0_img2 | 0.2 | 0.7 | 0.6 | 0.2 | 0.3 | 0.5 | 2.3 | 0.7 | 4 | 1.7 | 0.3 | 1 | 0.3 | 0 | 4 | 0 | 0 | 0 |
| Empty Bg Density Range Sigxy 1 SigZ 2/Poiss_30spots_bg_200_2_I_300_0_img0 | 0.2 | 0.7 | 0.6 | 0.2 | 0.2 | 0.5 | 0 | 0 | 0 | 0 | 0 | 0 | 0 | 0 | 3.3 | 0 | 0 | 0 |
| Empty Bg Density Range Sigxy 1 SigZ 2/Poiss_30spots_bg_200_2_I_300_0_img1 | 0.1 | 0.6 | 0.6 | 0.1 | 0.2 | 0.5 | 0 | 0 | 0 | 0 | 0 | 0 | 0 | 0 | 3.3 | 0 | 0 | 0 |
| Empty Bg Density Range Sigxy 1 SigZ 2/Poiss_30spots_bg_200_2_I_300_0_img2 | 0.1 | 0.6 | 0.5 | 0.1 | 0.2 | 0.5 | 0 | 0 | 0 | 0 | 0 | 0 | 0 | 0 | 13 | 0 | 0 | 0 |
| Empty Bg Density Range Sigxy 1pt5 SigZ 2/Poiss_300spots_bg_200_2_I_300_0_img0 | 0.3 | 0.8 | 0.6 | 0.3 | 0.4 | 0.6 | 4.3 | 2.3 | 3.7 | 2.3 | 2 | 1.7 | 0.3 | 0.3 | 4 | 0.3 | 0 | 0.3 |
| Empty Bg Density Range Sigxy 1pt5 SigZ 2/Poiss_300spots_bg_200_2_I_300_0_img1 | 0.3 | 0.9 | 0.6 | 0.3 | 0.5 | 0.8 | 4 | 2.7 | 8 | 3.3 | 3 | 4.3 | 0 | 0.3 | 5 | 4.3 | 0 | 3.3 |
| Empty Bg Density Range Sigxy 1pt5 SigZ 2/Poiss_300spots_bg_200_2_I_300_0_img2 | 0.2 | 0.8 | 0.6 | 0.3 | 0.4 | 0.7 | 3.7 | 2.7 | 4.3 | 3 | 2.7 | 3.3 | 0 | 0 | 5.3 | 2.3 | 0 | 1.3 |
| Empty Bg Density Range Sigxy 1pt5 SigZ 2/Poiss_30spots_bg_200_2_I_300_0_img0 | 0.2 | 0.9 | 0.7 | 0.2 | 0.4 | 0.5 | 0 | 0 | 3.3 | 0 | 0 | 0 | 0 | 0 | 10 | 0 | 0 | 0 |
| Empty Bg Density Range Sigxy 1pt5 SigZ 2/Poiss_30spots_bg_200_2_I_300_0_img1 | 0.2 | 0.9 | 0.6 | 0.2 | 0.4 | 0.5 | 0 | 0 | 0 | 0 | 0 | 0 | 0 | 0 | 6.7 | 0 | 0 | 0 |
| Empty Bg Density Range Sigxy 1pt5 SigZ 2/Poiss_30spots_bg_200_2_I_300_0_img2 | 0.2 | 0.8 | 0.5 | 0.2 | 0.3 | 0.6 | 0 | 0 | 0 | 0 | 0 | 0 | 0 | 0 | 10 | 0 | 0 | 0 |
| Empty Bg Density Range Sigxy 2 SigZ 2/Poiss_300spots_bg_200_2_I_300_0_img0 | 0.4 | 1 | 0.8 | 0.5 | 0.7 | 0.8 | 10 | 16 | 13 | 9.3 | 6 | 8.7 | 2 | 4.3 | 4.7 | 0 | 4 | 7.7 |
| Empty Bg Density Range Sigxy 2 SigZ 2/Poiss_300spots_bg_200_2_I_300_0_img1 | 0.4 | 1 | 0.8 | 0.5 | 0.7 | 0.8 | 13 | 17 | 14 | 13 | 7.7 | 9 | 1.7 | 4 | 7.3 | 0.7 | 5 | 9 |
| Empty Bg Density Range Sigxy 2 SigZ 2/Poiss_30spots_bg_200_2_I_300_0_img0 | 0.3 | 1.1 | 0.7 | 0.5 | 0.6 | 0.6 | 0 | 0 | 0 | 0 | 0 | 0 | 0 | 0 | 3.3 | 0 | 0 | 0 |
| Empty Bg Density Range Sigxy 2 SigZ 2/Poiss_30spots_bg_200_2_I_300_0_img1 | 0.2 | 1 | 0.8 | 0.4 | 0.7 | 0.9 | 10 | 3.3 | 0 | 3.3 | 3.3 | 0 | 3.3 | 0 | 17 | 0 | 3.3 | 3.3 |
| Empty Bg Density Range Sigxy 2 SigZ 2/Poiss_30spots_bg_200_2_I_300_0_img2 | 0.3 | 1 | 0.8 | 0.5 | 0.6 | 0.6 | 0 | 0 | 0 | 6.7 | 0 | 0 | 0 | 0 | 23 | 3.3 | 0 | 0 |
| Empty Bg SNR Range Sigxy 1 SigZ 2/Poiss_30spots_bg_200_1_I_300_0_img0 | 0.1 | 0.6 | 0.6 | 0.1 | 0.2 | 0.5 | 3.3 | 0 | 3.3 | 0 | 0 | 0 | 0 | 0 | 3.3 | 0 | 0 | 0 |
| Empty Bg SNR Range Sigxy 1 SigZ 2/Poiss_30spots_bg_200_1_I_300_0_img1 | 0.1 | 0.9 | 0.7 | 0.1 | 0.2 | 0.5 | 0 | 0 | 0 | 0 | 0 | 0 | 0 | 0 | 0 | 0 | 0 | 0 |
| Empty Bg SNR Range Sigxy 1 SigZ 2/Poiss_30spots_bg_200_1_I_300_0_img2 | 0.1 | 0.9 | 0.7 | 0.1 | 0.2 | 0.5 | 0 | 0 | 0 | 0 | 0 | 0 | 0 | 0 | 0 | 0 | 0 | 0 |
| Empty Bg SNR Range Sigxy 1 SigZ 2/Poiss_30spots_bg_200_2_I_300_0_img0 | 0.1 | 0.6 | 0.5 | 0.1 | 0.2 | 0.5 | 0 | 0 | 0 | 0 | 0 | 0 | 0 | 0 | 3.3 | 0 | 0 | 0 |
| Empty Bg SNR Range Sigxy 1 SigZ 2/Poiss_30spots_bg_200_2_I_300_0_img1 | 0.1 | 0.7 | 0.6 | 0.2 | 0.3 | 0.5 | 0 | 0 | 0 | 0 | 0 | 0 | 0 | 0 | 10 | 0 | 0 | 0 |
| Empty Bg SNR Range Sigxy 1 SigZ 2/Poiss_30spots_bg_200_2_I_300_0_img2 | 0.1 | 0.9 | 0.6 | 0.1 | 0.2 | 0.5 | 0 | 0 | 0 | 0 | 0 | 0 | 0 | 0 | 6.7 | 0 | 0 | 0 |
| Empty Bg SNR Range Sigxy 1 SigZ 2/Poiss_30spots_bg_200_4_I_300_0_img0 | 0.2 | 1 | 0.8 | 0.2 | 0.3 | 0.6 | 0 | 0 | 0 | 0 | 0 | 0 | 0 | 0 | 10 | 0 | 0 | 0 |
| Empty Bg SNR Range Sigxy 1 SigZ 2/Poiss_30spots_bg_200_4_I_300_0_img1 | 0.2 | 0.7 | 0.6 | 0.2 | 0.3 | 0.5 | 0 | 0 | 0 | 0 | 0 | 0 | 0 | 0 | 6.7 | 0 | 0 | 0 |
| Empty Bg SNR Range Sigxy 1 SigZ 2/Poiss_30spots_bg_200_4_I_300_0_img2 | 0.2 | 0.8 | 0.6 | 0.2 | 0.3 | 0.5 | 0 | 0 | 0 | 0 | 0 | 0 | 0 | 0 | 3.3 | 0 | 0 | 0 |
| Empty Bg SNR Range Sigxy 1 SigZ 2/Poiss_30spots_bg_200_8_I_300_0_img0 | 0.3 | 1 | 0.8 | 0.3 | 0.7 | 0.6 | 27 | 30 | 20 | 6.7 | 0 | 0 | 0 | 20 | 6.7 | 0 | 0 | 0 |
| Empty Bg SNR Range Sigxy 1 SigZ 2/Poiss_30spots_bg_200_8_I_300_0_img1 | 0.4 | 1.1 | 0.8 | 0.4 | 0.7 | 0.6 | 13 | 33 | 10 | 10 | 3.3 | 3.3 | 13 | 20 | 6.7 | 0 | 3.3 | 3.3 |
| Empty Bg SNR Range Sigxy 1 SigZ 2/Poiss_30spots_bg_200_8_I_300_0_img2 | 0.4 | 1 | 0.6 | 0.4 | 0.7 | 0.7 | 37 | 50 | 17 | 6.7 | 6.7 | 6.7 | 0 | 20 | 6.7 | 10 | 6.7 | 0 |
| Empty Bg SNR Range Sigxy 1pt5 SigZ 2/Poiss_30spots_bg_200_1_I_300_0_img0 | 0.2 | 0.8 | 0.6 | 0.2 | 0.3 | 0.7 | 3.3 | 0 | 6.7 | 3.3 | 3.3 | 3.3 | 0 | 0 | 6.7 | 0 | 3.3 | 0 |
| Empty Bg SNR Range Sigxy 1pt5 SigZ 2/Poiss_30spots_bg_200_1_I_300_0_img1 | 0.2 | 0.8 | 0.6 | 0.2 | 0.3 | 0.5 | 0 | 0 | 0 | 0 | 0 | 0 | 0 | 0 | 6.7 | 0 | 0 | 0 |
| Empty Bg SNR Range Sigxy 1pt5 SigZ 2/Poiss_30spots_bg_200_1_I_300_0_img2 | 0.2 | 1 | 0.6 | 0.2 | 0.3 | 0.5 | 0 | 0 | 3.3 | 0 | 0 | 0 | 0 | 0 | 17 | 0 | 0 | 0 |
| Empty Bg SNR Range Sigxy 1pt5 SigZ 2/Poiss_30spots_bg_200_2_I_300_0_img0 | 0.2 | 0.7 | 0.5 | 0.2 | 0.4 | 0.5 | 0 | 0 | 0 | 0 | 0 | 0 | 0 | 0 | 3.3 | 0 | 0 | 0 |
| Empty Bg SNR Range Sigxy 1pt5 SigZ 2/Poiss_30spots_bg_200_2_I_300_0_img1 | 0.2 | 0.9 | 0.6 | 0.2 | 0.3 | 0.6 | 0 | 0 | 0 | 0 | 0 | 0 | 0 | 0 | 13 | 0 | 0 | 0 |
| Empty Bg SNR Range Sigxy 1pt5 SigZ 2/Poiss_30spots_bg_200_2_I_300_0_img2 | 0.2 | 0.9 | 0.5 | 0.2 | 0.3 | 0.5 | 0 | 0 | 0 | 0 | 0 | 0 | 0 | 0 | 10 | 0 | 0 | 0 |
| Empty Bg SNR Range Sigxy 1pt5 SigZ 2/Poiss_30spots_bg_200_4_I_300_0_img0 | 0.3 | 1 | 0.7 | 0.3 | 0.5 | 0.6 | 6.7 | 0 | 0 | 0 | 0 | 0 | 0 | 0 | 6.7 | 0 | 0 | 0 |
| Empty Bg SNR Range Sigxy 1pt5 SigZ 2/Poiss_30spots_bg_200_4_I_300_0_img1 | 0.3 | 0.9 | 0.8 | 0.3 | 0.5 | 0.6 | 3.3 | 0 | 3.3 | 0 | 0 | 0 | 0 | 0 | 17 | 0 | 0 | 0 |
| Empty Bg SNR Range Sigxy 1pt5 SigZ 2/Poiss_30spots_bg_200_4_I_300_0_img2 | 0.3 | 0.9 | 0.8 | 0.3 | 0.5 | 0.6 | 13 | 0 | 6.7 | 0 | 0 | 0 | 3.3 | 0 | 13 | 0 | 0 | 0 |
| Empty Bg SNR Range Sigxy 2 SigZ 2/Poiss_30spots_bg_200_1_I_300_0_img0 | 0.2 | 1 | 0.6 | 0.4 | 0.5 | 0.7 | 0 | 0 | 0 | 0 | 0 | 0 | 0 | 0 | 10 | 0 | 0 | 0 |
| Empty Bg SNR Range Sigxy 2 SigZ 2/Poiss_30spots_bg_200_1_I_300_0_img1 | 0.2 | 1 | 0.7 | 0.4 | 0.4 | 0.5 | 0 | 0 | 0 | 3.3 | 0 | 0 | 0 | 0 | 13 | 3.3 | 0 | 0 |
| Empty Bg SNR Range Sigxy 2 SigZ 2/Poiss_30spots_bg_200_1_I_300_0_img2 | 0.2 | 0.9 | 0.6 | 0.4 | 0.5 | 0.5 | 0 | 0 | 0 | 0 | 0 | 0 | 0 | 0 | 10 | 0 | 0 | 0 |
| Empty Bg SNR Range Sigxy 2 SigZ 2/Poiss_30spots_bg_200_2_I_300_0_img0 | 0.2 | 1 | 0.6 | 0.5 | 0.5 | 0.5 | 0 | 0 | 3.3 | 10 | 0 | 0 | 0 | 0 | 17 | 0 | 0 | 0 |
| Empty Bg SNR Range Sigxy 2 SigZ 2/Poiss_30spots_bg_200_2_I_300_0_img1 | 0.3 | 1 | 0.8 | 0.5 | 0.6 | 0.6 | 0 | 0 | 0 | 6.7 | 0 | 0 | 0 | 0 | 10 | 0 | 0 | 0 |
| Empty Bg SNR Range Sigxy 2 SigZ 2/Poiss_30spots_bg_200_2_I_300_0_img2 | 0.2 | 1 | 0.7 | 0.5 | 0.5 | 0.6 | 0 | 0 | 3.3 | 10 | 0 | 0 | 0 | 0 | 23 | 3.3 | 0 | 0 |
| Infinite SNR Density Range Sigxy 1pt35 SigZ 2/Poiss_300spots_bg_200_0_I_10000_0_img0 | 0.1 | 0.6 | 0.4 | 0.1 | 0.1 | 0.5 | 2 | 1.3 | 4 | 1 | 1 | 1 | 0 | 0 | 3.7 | 0 | 0 | 0 |
| Infinite SNR Density Range Sigxy 1pt35 SigZ 2/Poiss_300spots_bg_200_0_I_10000_0_img1 | 0.1 | 0.6 | 0.4 | 0.1 | 0.1 | 0.5 | 4 | 2.3 | 7.3 | 2.3 | 2 | 2 | 0 | 0 | 3 | 0 | 0 | 0 |
| Infinite SNR Density Range Sigxy 1pt35 SigZ 2/Poiss_300spots_bg_200_0_I_10000_0_img2 | 0.2 | 0.6 | 0.4 | 0.1 | 0.1 | 0.5 | 4.7 | 3 | 5.7 | 2.3 | 1.7 | 2.3 | 0.3 | 0 | 3.7 | 0 | 0 | 0 |
| Infinite SNR Density Range Sigxy 1pt35 SigZ 2/Poiss_30spots_bg_200_0_I_10000_0_img0 | 0 | 0.6 | 1 | 0 | 0 | 0.5 | 0 | 0 | 0 | 0 | 0 | 0 | 0 | 0 | 3.3 | 0 | 0 | 0 |
| Infinite SNR Density Range Sigxy 1pt35 SigZ 2/Poiss_30spots_bg_200_0_I_10000_0_img1 | 0 | 0.6 | 1 | 0 | 0 | 0.5 | 0 | 0 | 0 | 0 | 0 | 0 | 0 | 0 | 3.3 | 0 | 0 | 0 |
| Infinite SNR Density Range Sigxy 1pt35 SigZ 2/Poiss_30spots_bg_200_0_I_10000_0_img2 | 0 | 0.6 | 0.9 | 0 | 0 | 0.5 | 0 | 0 | 0 | 0 | 0 | 0 | 0 | 0 | 10 | 0 | 0 | 0 |
| **MEAN** | **0.2** | **0.8** | **0.7** | **0.25** | **0.36** | **0.6** | **3.4** | **3.3** | **3.1** | **2.2** | **0.9** | **1** | **0.5** | **1.4** | **7.9** | **0.6** | **0.5** | **0.6** |

**Supplementary Table 2**: Comparison of execution time of real 3D datasets. The tool's parameters were set for each image so that the number of detected spots was comparable across tools (excluding deepBlink, as no parameters could be set). The tools compared are AIRLOCALIZE (Air), Big-FISH (BF), deepBlink (DB), FISH-quant (FQ), RS-FISH (RS), and Starfish (SF).

| | Number of Detected Spots | | | | | | Computation Time | | | | | |
|---|---|---|---|---|---|---|---|---|---|---|---|---|
| **method** | **Air** | **BF** | **DB** | **FQ** | **RS** | **SF** | **Air** | **BF** | **DB** | **FQ** | **RS** | **SF** |
| **name** | | | | | | | | | | | | |
| C0_CB428_22_cropped_4065 | 1580 | 1695 | 319 | 1597 | 1649 | 1648 | 5.52 | 3.36 | 4.56 | 7.5 | 2.64 | 4.72 |
| C0_N2_1093_cropped_3801 | 1098 | 1210 | 53 | 1043 | 1124 | 1167 | 3.57 | 2.88 | 5.1 | 6.4 | 0.74 | 4.15 |
| C0_N2_1121_cropped_3785 | 972 | 999 | 51 | 979 | 969 | 989 | 3.31 | 2.511 | 3.69 | 6.9 | 0.71 | 3.22 |
| C0_N2_1449_cropped_3615 | 311 | 293 | 332 | 348 | 283 | 302 | 1.14 | 1.89 | 3.59 | 4.2 | 0.51 | 2.2 |
| C0_N2_1639_cropped_3974 | 600 | 571 | 35 | 569 | 562 | 599 | 2.02 | 2.19 | 3.42 | 4.5 | 0.52 | 2.65 |
| C0_N2_702_cropped_1620 | 2160 | 2137 | 213 | 2049 | 2119 | 1897 | 7.85 | 2.07 | 3.52 | 8.8 | 1.24 | 3.83 |
| C1_CB428_22_cropped_4065 | 46 | 29 | 4 | 77 | 31 | 34 | 0.35 | 3.59 | 4.28 | 3.4 | 0.4 | 4.11 |
| C1_N2_1449_cropped_3615 | 96 | 50 | 13 | 93 | 55 | 0 | 0.51 | 1.93 | 3.35 | 2.8 | 0.26 | 2.08 |
| C2_CB428_22_cropped_4065 | 2130 | 1707 | 106 | 2122 | 1778 | 1712 | 8.75 | 3.52 | 4.29 | 9.6 | 1.01 | 5.53 |
| C2_N2_1121_cropped_3785 | 27 | 26 | 3 | 58 | 30 | 36 | 0.24 | 2.46 | 3.41 | 3.4 | 0.28 | 2.69 |
| C2_N2_1449_cropped_3615 | 364 | 305 | 56 | 320 | 329 | 310 | 1.23 | 1.95 | 3.43 | 4.2 | 0.33 | 2.53 |
| C2_N2_702_cropped_1620 | 273 | 183 | 4 | 213 | 182 | 224 | 0.93 | 2.12 | 3.5 | 3.3 | 0.3 | 2.51 |
| C2_RNAi_dpy27_1369_cropped_5751 | 45 | 154 | 1 | 90 | 151 | 129 | 0.5 | 3.41 | 4.13 | 2 | 0.42 | 3.68 |
| **MEAN** | | | | | | | **2.76** | **2.61** | **3.87** | **5.154** | **0.72** | **3.377** |

**Supplementary Table 3:** Information on datasets used for this study.

| **dataset** | **organism** | **short description** | **reference** |
|---|---|---|---|
| Fig 1c | *C. elegans* | *C. elegans* larvae labeled with smFISH for *lea-1* mRNA. | This study |
| Fig 1d | mESC | Mouse embryonic stem cell labeled with DAPI (blue) and smFISH for cdx-2 mRNA (white). | This study |
| Fig 1e | *Drosophila* | *Drosophila* brain labeled with smFISH for Pura mRNA. | This study |
| Fig 1f | PGP1f cells | Images correspond to four channels of a single round of OligoFISSEQ labeling with the 36plex-5K oligopaint library (Nguyen et al.[9], Fig. 2) | Nguyen et al.[9] |
| Fig 1g | *C. elegans* | One example image of a *C. elegans* embryo labeled with smFISH for *mdh-1* mRNA of a dataset with 4010 mixed stage embryos. | This study |
| Fig 1h, Supplementary Video 1 | mouse | Expansion-Assisted Iterative Fluorescence In Situ Hybridization (EASI-FISH) spots for the gene Map1b detected in a tissue section of the lateral hypothalamic area. | Wang et al.[10] |
| Fig 2 a-c, Supplementary Table 1 | simulated | Simulated data of diffraction-limited spots with varying degrees of background noise. | This study |
| Fig 2d, Supplementary Table 2 | *C. elegans* | 13 *C. elegans* embryos at different developmental stages labeled with smFISH for different mRNAs. | This study |

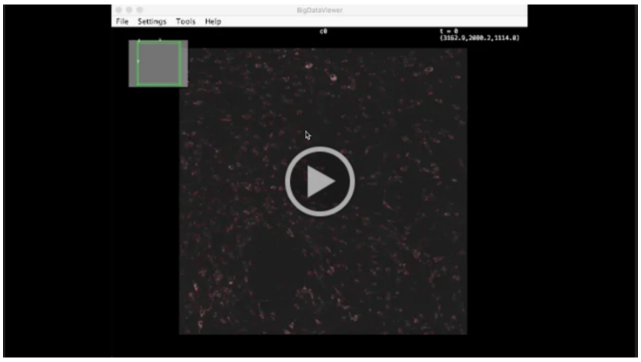

https://drive.google.com/file/d/1vUS5tih5NJlniEGM4Cr-b5AjBy8f_wtk/view?usp=sharing

**Supplementary Video 1** Fly-through video of a large lightsheet dataset with RS-FISH detected spots using BigDataViewer. A 148 GB lightsheet dataset of a tissue section of the lateral hypothalamic area labeled using EASI-FISH (grayscale) and spots detected using RS-FISH (red).

### Derivation of the three-dimensional Radial Symmetry Localization

**(and obvious extension to the n-dimensional case)**

Our extension of Radial Symmetry from two dimensions (2D) to three dimensions (3D) requires adjustments of gradient computation and is based on a new derivation that can scale to the n-dimensional case. Although we only show the 3D case here, we suggest how to straightforward extend these concepts to higher dimensions. General concepts are illustrated in **Supplementary Figure 1**.

First, image gradients $\nabla I(p)$ are computed, which are located on a half-pixel offset grid (**Supplementary Figure. 1a,b**). Parthasarathy[1] suggested using the Roberts cross operator, which, however, cannot be straightforward extended to 3D. Instead, we use separable computation of image gradients for the 2D case:

$$\nabla I(p) = \begin{bmatrix} \frac{\partial I}{\partial x}(p) \\ \frac{\partial I}{\partial y}(p) \end{bmatrix} = \begin{bmatrix} \frac{I(p_1)+I(p_3)-I(p_0)-I(p_2)}{2} \\ \frac{I(p_2)+I(p_3)-I(p_0)-I(p_1)}{2} \end{bmatrix}$$

and the 3D case:

$$\nabla I(p) = \begin{bmatrix} \frac{\partial I}{\partial x}(p) \\ \frac{\partial I}{\partial y}(p) \\ \frac{\partial I}{\partial z}(p) \end{bmatrix} = \begin{bmatrix} \frac{I(p_1)+I(p_3)+I(p_5)+I(p_7)-I(p_0)-I(p_2)-I(p_4)-I(p_6)}{4} \\ \frac{I(p_2)+I(p_3)+I(p_6)+I(p_7)-I(p_0)-I(p_1)-I(p_4)-I(p_5)}{4} \\ \frac{I(p_4)+I(p_5)+I(p_6)+I(p_7)-I(p_0)-I(p_1)-I(p_2)-I(p_3)}{4} \end{bmatrix}$$

Due to the separable nature of the computation, it can be easily extended to higher dimensions.

Using these image gradients, it is possible to compute their 3D intersection point $p_c = (x_c, y_c, z_c)$. In contrast to the original 2D radial symmetry, we derive the formula for computing the shortest distance $d_k$ between $p_c$ and the vector $\overrightarrow{v_k}$ defined by the location $p_k = (x_k, y_k, z_k)$ and the gradient $\nabla I(p_k)$ using vector algebra as it extends to higher dimensions (**Supplementary Figure 1c**). Therefore, we compute the dot product between the vector $\overrightarrow{v_c} = p_c - p_k$ and $\overrightarrow{v_k} = p_k + \dot{\nabla} I(p_k)$ which projects $\overrightarrow{v_c}$ onto $\overrightarrow{v_k}$ computing the helper distance $e_k$:

$$e_k = |\overrightarrow{v_c}| \cdot \frac{\overrightarrow{v_k}}{|\overrightarrow{v_k}|}$$

From that, one can directly compute

$$d_k = |p_c - e_k \overrightarrow{v_k}|$$

Replacing the actual coordinates and reformulating the above equation yields for the 3D case:

$$\Delta x = x_k - x_c$$
$$\Delta y = y_k - y_c$$
$$\Delta z = z_k - z_c$$
$$\alpha = a_k * \Delta x + b_k * \Delta y + c_k * \Delta z$$
$$d_k^2 = \Delta x^2 + \Delta y^2 + \Delta z^2 - \frac{\alpha^2}{a_k^2 + b_k^2 + c_k^2}$$

for which the pattern for extension to the n-dimensional case becomes obvious when comparing it to the 2D case:

$$\Delta x = x_k - x_c$$
$$\Delta y = y_k - y_c$$
$$\alpha = a_k * \Delta x + b_k * \Delta y$$
$$d_k^2 = \Delta x^2 + \Delta y^2 - \frac{\alpha^2}{a_k^2 + b_k^2}$$

We then derived the least-squares fit for identifying the common intersection point $p^c = (x_c, y_c, z_c)$ of all gradients by minimizing (**Supplementary Figure 1d**):

$$\chi^2 \equiv \sum_k d_k^2 w_k$$

where $w_k$ denotes optional weights for each gradient $\nabla I(p^k)$. Derivations for the 3D case:

$$\frac{\partial \chi^2}{\partial x_c} = 0, \qquad \frac{\partial \chi^2}{\partial y_c} = 0, \qquad \frac{\partial \chi^2}{\partial z_c} = 0$$

yields a set of linear equations that can be expressed in matrix form $\Delta * p_c = \Theta$

$$m_k^2 = a_k^2 + b_k^2 + c_k^2$$

$$\begin{bmatrix} \sum_k 1 - \frac{a_k^2}{m_k^2} & \sum_k -\frac{a_k b_k}{m_k^2} & \sum_k -\frac{a_k c_k}{m_k^2} \\ \sum_k -\frac{a_k b_k}{m_k^2} & \sum_k 1 - \frac{b_k^2}{m_k^2} & \sum_k -\frac{b_k c_k}{m_k^2} \\ \sum_k -\frac{a_k c_k}{m_k^2} & \sum_k -\frac{b_k c_k}{m_k^2} & \sum_k 1 - \frac{c_k^2}{m_k^2} \end{bmatrix} * \begin{bmatrix} x_c \\ y_c \\ z_c \end{bmatrix} = \begin{bmatrix} \sum_k x_k \left(1 - \frac{a_k^2}{m_k^2}\right) - \left(\frac{a_k b_k y_k}{m_k^2}\right) - \left(\frac{a_k c_k z_k}{m_k^2}\right) \\ \sum_k y_k \left(1 - \frac{b_k^2}{m_k^2}\right) - \left(\frac{a_k b_k x_k}{m_k^2}\right) - \left(\frac{b_k c_k z_k}{m_k^2}\right) \\ \sum_k z_k \left(1 - \frac{c_k^2}{m_k^2}\right) - \left(\frac{a_k c_k x_k}{m_k^2}\right) - \left(\frac{b_k c_k y_k}{m_k^2}\right) \end{bmatrix}$$

Interestingly, the pattern for extension to the n-dimensional case becomes again obvious when comparing it to the 2D case:

$$m_k^2 = a_k^2 + b_k^2$$

$$\begin{bmatrix} \sum_k 1 - \frac{a_k^2}{m_k^2} & \sum_k -\frac{a_k b_k}{m_k^2} \\ \sum_k -\frac{a_k b_k}{m_k^2} & \sum_k 1 - \frac{b_k^2}{m_k^2} \end{bmatrix} * \begin{bmatrix} x_c \\ y_c \end{bmatrix} = \begin{bmatrix} \sum_k x_k \left(1 - \frac{a_k^2}{m_k^2}\right) - \left(\frac{a_k b_k y_k}{m_k^2}\right) \\ \sum_k y_k \left(1 - \frac{b_k^2}{m_k^2}\right) - \left(\frac{a_k b_k x_k}{m_k^2}\right) \end{bmatrix}$$

These systems of linear equations can be solved by inverting Δ:

$$p_c = \Delta^{-1} * \Theta$$

For the 2D and 3D case we invert Δ using closed-form solutions. Inverting higher-dimensional matrices can be for example achieved by computing the pseudo-inverse using Singular Value Decomposition[2].

### Derivation for supporting axis-aligned ellipsoid (non-radial) objects

Radial Symmetry is, as the name suggests, designed for round objects where all gradients intersect exactly at the center point. However, it is desirable to support ellipsoid objects in order to support for example anisotropic datasets common in microscopy.

In microscopy, the pixel size in lateral ($xy$) and axial dimension ($z$) are not identical. The $xy$ dimension is typically acquired with equal pixel size in $x$ and $y$ using a camera or scanning device collecting light from an objective, while the step size in $z$ is usually defined by a motor that moves the sample or the objective. Typically, the pixel size in $z$ is bigger by a factor of 2-10. At the same time, the point spread function (PSF) of a microscopic system is also much smaller in $xy$ compared to $z$. Although these two effects kind of balance each other, imaged single-molecule spots do not appear as round but ellipsoid objects in 3D.

We therefore derived a version of radial symmetry that supports axis-aligned, ellipsoid objects. We define the scaling for each gradient location as

$$\begin{pmatrix} x_k \\ u_k \\ v_k \end{pmatrix}^T \circ \begin{pmatrix} 1 \\ \alpha \\ \beta \end{pmatrix}^T = \begin{pmatrix} x_k \\ y_k \\ z_k \end{pmatrix}^T$$

where $q_k = (x_k, u_k, v_k)$ describes anisotropic coordinates in the input image, $s = (1, \alpha, \beta)$ describes the global known scale factors (anisotropy factors), $p_k = (x_k, y_k, z_k)$ describes the isotropic space coordinates in which radial symmetry will be computed, and $\circ$ denotes element-wise multiplication (or Hadamard product). Mapping the gradients extracted from input images $\nabla I(q_k)$ at coordinates $q_k$ into isotropic space with coordinates $p_k$ yields:

$$\nabla I(p) = \begin{bmatrix} \frac{\partial I}{\partial x}(p) \\ \frac{\partial I}{\partial y}(p) \\ \frac{\partial I}{\partial z}(p) \end{bmatrix} = \begin{bmatrix} \frac{\partial I}{\partial x}(q) \\ \frac{\partial I}{\partial u}\frac{\partial u}{\partial y}(q) \\ \frac{\partial I}{\partial v}\frac{\partial v}{\partial z}(q) \end{bmatrix} = \begin{bmatrix} 1 \\ {}^{1}\!/_{\alpha} \\ {}^{1}\!/_{\beta} \end{bmatrix} \circ \begin{bmatrix} \frac{\partial I}{\partial x}(q) \\ \frac{\partial I}{\partial u}(q) \\ \frac{\partial I}{\partial v}(q) \end{bmatrix} = \begin{bmatrix} 1 \\ {}^{1}\!/_{\alpha} \\ {}^{1}\!/_{\beta} \end{bmatrix} \circ \nabla I(q)$$

In summary, the anisotropic coordinates $q_k$ need to be scaled with the global scale vector $s$, while gradients extracted from the anisotropic input image $\nabla I(q_k)$ need to be scaled with the inverse of the global scale in order to compute radial symmetry on anisotropic datasets. Again, scaling to n-dimensional datasets is straightforward. Importantly, the resulting center point $p_c = (x_c, y_c, z_c)$ will be in isotropic space and might need to be scaled back into the anisotropic space.

### Computing the global scale factor

For single-molecule detections, the global scale vector depends mainly on the pixel size and the PSF. For practical reasons we only support computation of the $z$ scale factor $\beta$ (called anisotropy factor) in the Fiji plugin. While assuming a scale of $\beta = 1.0$ might yield reasonable results due to the fact that objects are usually roughly spheres, we highly recommend computing it for a specific set of microscope settings that are usually held constant across many experiments (same objective, same z stepping, same camera, same magnification).

The RS-FISH plugin offers two possibilities for computing the anisotropy factor. First, points can be interactively identified using Difference-of-Gaussian and a 3D Gaussian is fitted to each detection, which directly computes the anisotropy. Second, RS- FISH can sequentially run RS-RANSAC on a range of anisotropy factors (typically $0.1 - 5.0$). The correct anisotropy factor will be the one that maximizes the number of pixels used for spot detection since this indicates that most gradients intersect in the center of each spot. Both ways usually yield very comparable results, which can also be easily confirmed by manual inspection of single spots in 3D.

### Accuracy benchmarking RS-FISH against commonly used tools

We compared RS-FISH's accuracy with the five most commonly used spot detection tools - FISH-quant (v3) [3], FISH-quant's python version - Big-FISH (0.5.0) [4], AIRLOCALIZE (1.6) [5], Starfish (0.2.2) [6], deepBlink (0.1.1) [7] (**Fig. 2, Supplementary Table 1**). The accuracy of spot detection was benchmarked using 50 simulated images, each of size 264x264x32 and containing either 30 or 300 simulated spots. For each tool, excluding deepBlink, we ran a grid search of the parameter space within a range of sensible values per parameter, running all value combinations between parameters. Parameter grid search was not run on deepBlink as it is an artificial neural network-based tool, however, we did run multiple configurations in the search for optimal results. Some of the benchmarked tools offer more than one spot detection pipeline. In such cases, we chose only one pipeline by following their documentation. Notably, this raises the possibility that superior accuracy and execution time were overlooked. This is especially relevant for Starfish, as Starfish is a tool for building image processing pipelines. The next section will detail the methods and grid search parameter value combinations (inclusive) for each tool.

All benchmarking scripts for all tools and instructions on how to reproduce are found here:
https://github.com/PreibischLab/RS-FISH/tree/master/documents/tool_comparison_for_paper
Instructions for installation and running the grid search are found in the README file in the link above.

**RS-FISH**
Pipeline: RANSAC
grid search parameters:

DoGSigma = 1-2.5, step size += 0.25
DoGthreshold = 0.001-0.13, step size *= 1.5
supportRadius: 2-4, step size +=1
inlierRatio: 0.0-0.3, step size += 0.1
maxError: 0.1-2.6 step size += 0.5
intensityThreshold: [0,100,150,200]

**FISH-quant**
Pipeline: Gaussian Fit
As FISH-quant is a MATLAB GUI application, optimal parameters were found manually for each image.

**Big-FISH**
Pipeline: Dense region decomposition
As Big-FISH calculates a default value for each parameter automatically, the grid search range for some parameters was derived from the default value.
grid search parameters:

sigmaZ: default + [-0.5,-0.25,0,0.25,0.5]
sigmaYX: default + [-0.5,-0.25,0,0.25,0.5]
threshold: value of index in threshold array, with indices relative to location of the default.
Relative to default indices: [-6,-3,-2,-1,0,1,2,3,6]
alpha: 0.5-0.8, step size += 0.1

beta: 0.8-1.2, step size += 0.1
Gamma: 4-6, step size += 0.5

**AIRLOCALIZE**
Pipeline: 3DMaskFull

grid search parameters:
Threshold = 4-13, step size += 1
minDistBetweenSpots = 1-3, step size += 1

**Starfish**
Pipeline: BlobDetector

grid search parameters:
Sigma: 2-5, step-size += 1
Threshold: 0.00004-0.000095, step size += 0.000001, and additional customized threshold values for images with insufficient accuracy.

**deepBlink**
Pipeline: deepBlink's pre-trained network "Particle"
DeepBlink is an artificial neural network-based tool for spot detection in 2D images, while all of our comparisons were performed on 3D images. However, since they offer a 3D adaptation solution and as it is currently the only notable tool that is based on deep learning, we decided to include it in the benchmarking analysis.
We have run multiple networks and found "particle.h5" to perform best on our simulated data (as well the embryo images used in the execution time benchmarking detailed in the next section). The network outperformed deepBlink's pre-trained network "smfish.h5". It also outperformed networks that we trained from scratch, one specific network for each simulated image, using all available simulated images of the same signal-to-noise ratio as a training set. We suspect that the small training set and the small pool of spots were insufficient for the training task.

**Analyzing grid search results**

A grid search approach was used to find well-performing parameter combinations of RS-FISH and the other tools across 50 ground-truth simulated images, 38 images containing 30 spots each and 12 images containing 300 spots each. The dataset consists of images with a varying signal-to-noise ratio.
The resulting list of spot detections of each parameter combination of each software was evaluated by comparing it to a list of ground truth points. Both lists are 3D coordinates (x,y,z), with the size of the list being the number of spots. First, a KD-Tree of all detected points was constructed. All ground truth points were queried against the KD-Tree. The ground truth list element with the smallest distance between itself and a detection was deleted from the ground truth list. The corresponding detection element was also deleted from the detection list. This process was repeated until there were either no elements left in the ground truth list, no elements left in the detection list, or the smallest distance between a

ground truth point and elements in the detection array was above a certain threshold (a distance of *2.0* was used).

Once an end condition was met, the performance of the parameter combination was assessed (**Supplementary Table 1**). The performance was measured using the length of the ground truth list after measurement (how many ground truth points have not been detected), and the length of the detection list after measurement (how many false detections have been made), and the average Euclidean distance between the detected spots and their associated ground truth.
Best performance of each software for each image was selected by choosing the parameter combination results that minimize the sum of missed detections (false negatives, FN) and false detections (false positives, FP). The optimal results would have a score of zero: zero missed detections (FN) and zero false detections (FP). In case multiple parameter combinations resulted in the same performance using this metric (sum of FP and FN), the average Euclidean distance between detections and their associated ground truth point was also taken into account, with the best results having the lowest mean Euclidean distance.
We compared the performance for each tool across ground truth simulated images containing different numbers of points and varying background noise (**Supplementary Table 1**).

### Computation time benchmarking

To compare computation time between RS-FISH, FISH-quant (v3)[3], Big-FISH (v5)[4], AIRLOCALIZE (1.6)[5], Starfish (0.2.2)[6], and deepBlink (0.1.1)[7], we performed several realistic smFISH localization runs on 13 3D images of *C. elegans* embryos (**Supplementary Table 2**). Image size varied between images with lateral size (XY) of 334-539 pixels and axial size (Z) of 81-101 pixels. An example is shown in **Fig. 1g**. Since execution times significantly depend on the number of spots being detected, we used settings that produce a comparable number of spots in all tools, excluding deepBlink. deepBlink spot detection was performed using its pre-trained artificial neural network "Particle", thus the approximate number of spots found was fixed. Notably, the difference in the results to other methods may be attributed to a mismatch between the chosen pre-trained network and the spot detection task on the embryo images, however, we did not have training data to train the network from scratch. Additionally, as deepBlink is based on networks to analyze 2D images, a sizable portion of each computation time that is reported was exhausted by the step that adapts the network's results from 2D to 3D, and while this step is offered as part of deepBlink's package, it is not part of the main pipeline.
In the metric of computation time, we found that RS-FISH significantly outperforms all compared tools.

Link to download embryo images:
bimsbstatic.mdc-berlin.de/akalin/preibisch/RSFISH_embryos.zip

The computation time benchmarks for all tools were run on a Dell Precision Tower 7910 workstation, Intel(R) Xeon(R) CPU E5-2640 v4 @ 2.40GHz. deepBlink spot detection was run with TensorFlow GPU on Nvidia Quadro P6000.

### Tutorial RS-FISH

#### Calculating Anisotropy Coefficient

Since the effective size of objects along the z-axis is usually different from the x- and y-axis of many images, it should be corrected to achieve a more accurate smFISH detection. To estimate your anisotropy coefficient, you can acquire a fluorescent bead image on the same microscope, using the same settings and equipment, or you can use the smFISH image directly.

Open the image with the beads or the smFISH detections and navigate to the 'Plugins' > 'RS-FISH' >'Tools'> 'Calculate Anisotropy Coefficient'.

You will see the dialog window in **Supplementary Figure 2a**. Make sure your bead image is selected in the **Image** drop-down menu. Next, you can choose between two **Detection methods: Gauss fit** or **Radial Symmetry.** If you have fewer detections, Gaussian fit might be the better choice; however, both methods usually provide reasonable results. It can even be beneficial to simply average the results of both approaches. The resulting number can be visually confirmed by turning the input image around its x or y-axis (Image > Stacks > Reslice > Top) as it simply describes the ratio of the size in Z versus XY. After you choose a detection method, two windows will open once you press **OK**.

In the **Adjust difference-of-gaussian values** window (**Supplementary Figure 2b**), you can choose **Sigma** and **Threshold** values to detect the majority of subpixel resolution spots.

Once you are done – press the **Done** button.

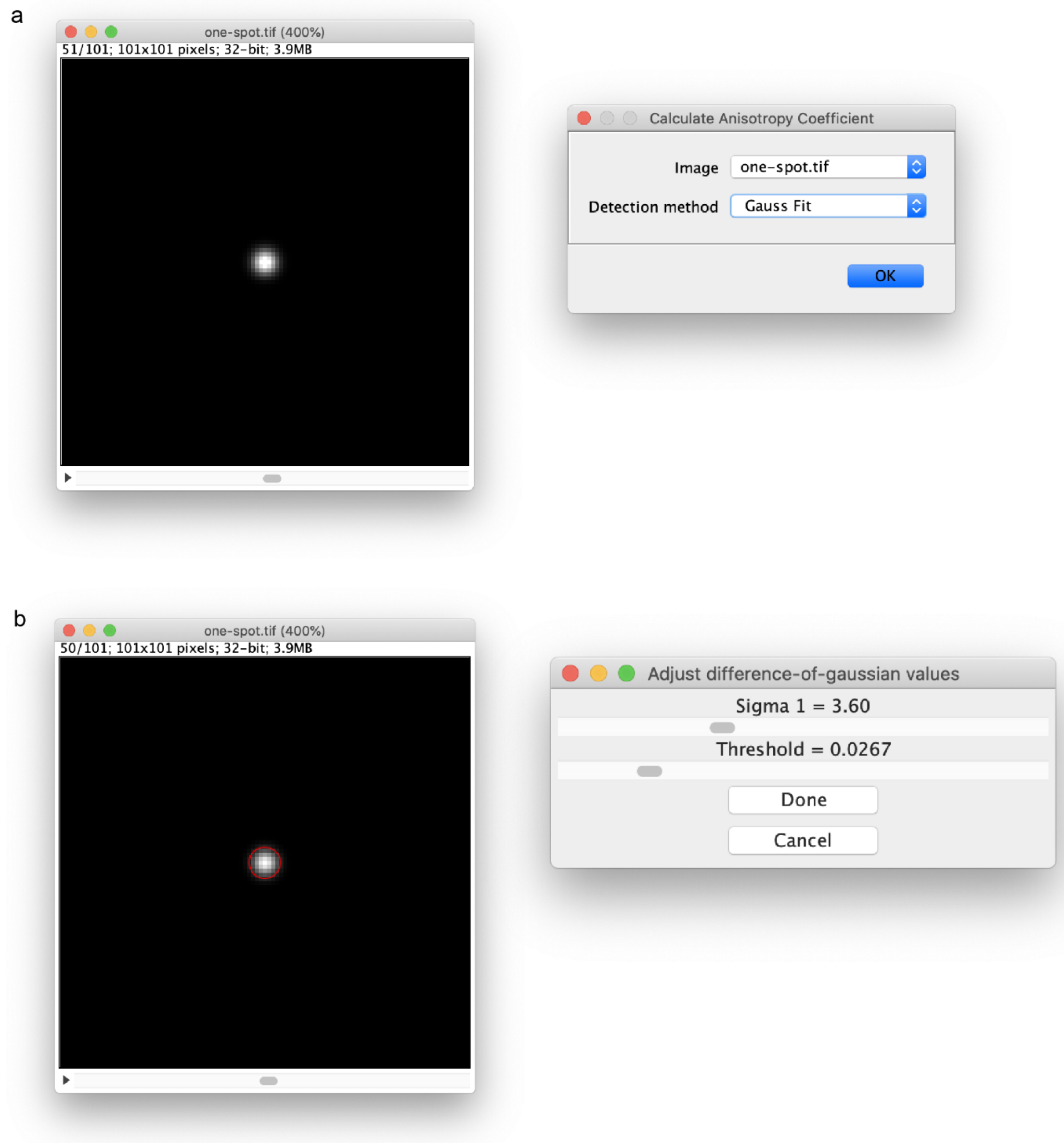


**Supplementary Figure 2**. Calculating the Anisotropy Coefficient of a fluorescent image**.** a) Selecting a bead or smFISH image for anisotropy coefficient detection. b) Adjusting the difference-of-gaussian values to detect single spots for anisotropy coefficient calculation.

Depending on the number of spots, the calculations might take some time as the Gaussian fit is slower, and the RS-RANSAC needs to iterate over a range of potential anisotropy coefficients. The program will calculate the corresponding anisotropy coefficient, which shows how we should squeeze the objects in the z-axis to make them look radially symmetric.

The Log window will show the corresponding anisotropy value, and it should be automatically transferred to the next step.

*Important: It is OK to skip this step if the objects are more or less round in 3D. The plugin will do a decent job even with the default value of the anisotropy coefficient. However, we advise performing this before the actual RS detection.*

#### Localizing Spots

The main RS-FISH plugin can be found under Plugins > RS-FISH > RS-FISH. There are two different modes of processing images: **interactive** and **advanced**. The Interactive method is used to adjust the parameters for further dataset processing or is the right choice if single images need to be processed. The interactive mode provides the visual feedback necessary to adjust the advanced automated processing parameters on large datasets.

#### Interactive mode

Open a 2D or 3D single-channel image for analysis and navigate to the 'Plugins' menu under 'RS-FISH' > 'RS-FISH'. A window will pop up, as depicted in **Supplementary Figure 3a**. Ensure that the correct image is chosen in the **Image** drop-down menu. Next, you choose the **mode** that you want to run RS-FISH in. For finding the best parameters or analyzing a small set of images, select the **Interactive Mode**.

**Anisotropy coefficient** defines how much the z-resolution differs from the x/y-resolution. The parameter would be set automatically if you ran the '**Calculate Anisotropy Coefficient**' plugin beforehand (Plugins > RS-FISH > Calculate Anisotropy Coefficient'). In general, 1.0 gives a good result if the spots are somewhat round in 3D. You can choose to use the same anisotropy coefficient for computing the Difference of Gaussian (DoG), which will lead to a more robust DoG detection for anisotropic spots.

There are various options for **Robust fitting Computation.**

- '**RANSAC**' defines if you want to use radial symmetry with robust outlier removal, it will identify all gradients within every local patch that supports the same center point (**Fig. 1**)
- '**No RANSAC**' for the use of radial symmetry without robust outlier removal, simply all gradients of a local spot will be used to compute the center point (classic RS)
- ‘**Multiconsensus RANSAC**’ will iteratively run robust outlier removal on each local patch until all sets of gradients are identified that support a center point. This allows RS-FISH to potentially find multiple points within each local patch that was identified using DoG detections.

The last option during this first step is to visually select the spots from an Intensity histogram in the **Visualization** section. This option is only available in the interactive mode. This option will allow you to choose the found smFISH spot by thresholding based on an intensity value. Once you are done with the settings, press the **OK** button.

In the second step, multiple windows will open based on your selection (**Supplementary Figure 3b**).

In the **Difference of Gaussian** window, you can adjust the parameters for the initial detection of the spots. The goal of this step is to minimize false detections. Adjust the **Sigma** and **Threshold** slider so that the red circles in the image detect as many single spots as possible. Try to slightly find more spots if you choose RANSAC; the RANSAC window allows additional restrictive settings. If you are working with a 3D stack, it helps to move through z while adjusting the parameters as the red circle appears only in the z-slices where the signal is the strongest. It can help to change the yellow preview box during this step. If the image is very large, it can help choose a smaller box to speed up the visualization (the detection will be performed in the whole image).

*Important: If you choose to run RANSAC robust fitting, do not click the Done button on the Difference of Gaussian window at this step; simply continue setting the parameters in the Adjust RANSAC values window.*

The **Adjust RANSAC values** dialog allows you to find the right setting for the robust outlier removal. The **Support Region Radius** defines the radius for each spot in which gradients are extracted for RS. You might want to play with this parameter. Sometimes it is helpful to increase the radius and decrease the **Inlier Ratio** at the same time. The **Inlier ratio** defines the ratio of the gradients (i.e., pixels) that have to support the RS center point (Simply speaking, the ratio of pixels should 'belong' to the current spot), given the **Max error** that defines maximally allowed error for RS fitting (see **Fig. 1**).

While moving the sliders, you will see the updates in both image windows. Firstly the **RANSAC preview** window displays the pixels used by RANSAC and the error values at each of the used pixels. The second window is the initial image window with the preview of the detections. Additionally to the red circles, the blue crosses indicate spots that were detected using RANSAC outlier removal. So the goal of this part is to find all spots with a red circle and a blue cross inside while not detecting noise or background.

The background removal step allows you to remove a non-planar background prior to computing the RS. It will try to estimate a plane using the intensity values of the edges of each local patch using any of the outlined methods. *Note: constant backgrounds do not need to be removed, only if strong gradients are present.*

Once the parameters are adjusted, hit any of the 'Done' buttons and wait a bit while the computations are performed.

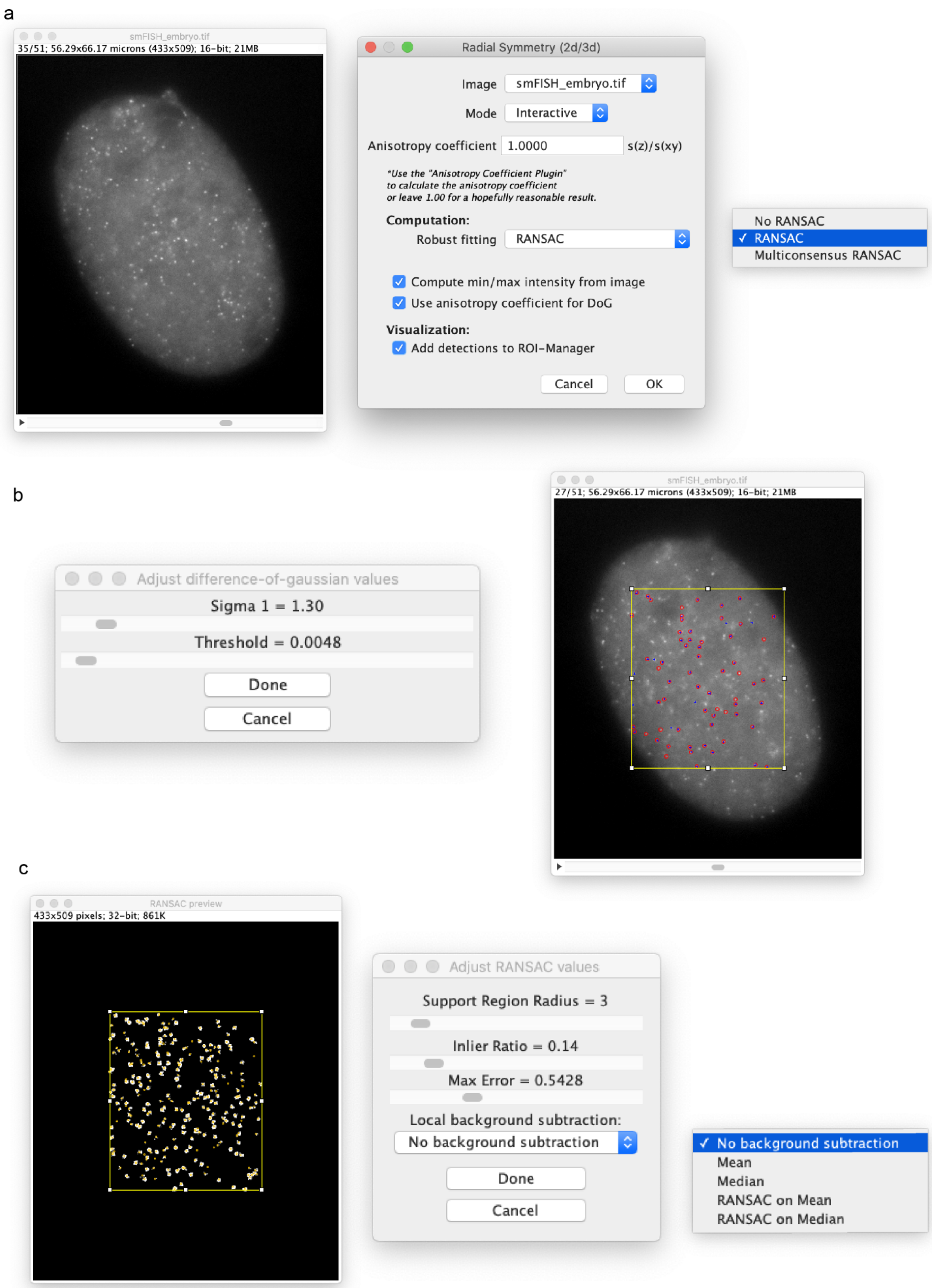


**Supplementary Figure 3:** The RS-FISH interactive mode for spot detection**.** a) With an open smFISH image, navigate to the RS-FISH plugin and choose the interactive mode. Choose the computation mode from the drop-down menu for fitting the spots. b) Adjust the difference-of-gaussian values using the sliders to detect single spots in your selected image within the yellow selection box. c) Adjust the RANSAC values for improved detection accuracy and correct an uneven background if necessary.

In the next window (**Supplementary Figure 4a**), you have the option of thresholding the detected spots based on their intensity. The **Intensity distribution** window displays all detected spots and their corresponding intensity value as a histogram. By clicking at an intensity value in the histogram, the blue thresholding bar can be adjusted. All spots that currently pass the thresholding are displayed in the image window and marked by a red circle. If you are satisfied with the selected spots, press the **OK** button and continue to the final results table.

The **Log** window (**Supplementary Figure 4b**) gives you a summary of all spots found at every step and the final number of detections. The **Results table** (**Supplementary Figure 4c**) contains the spot coordinates, time, channel, and intensity values in the corresponding columns. You can save the results and use them in the **Show Detections** plugin to visualize all found spots' locations.

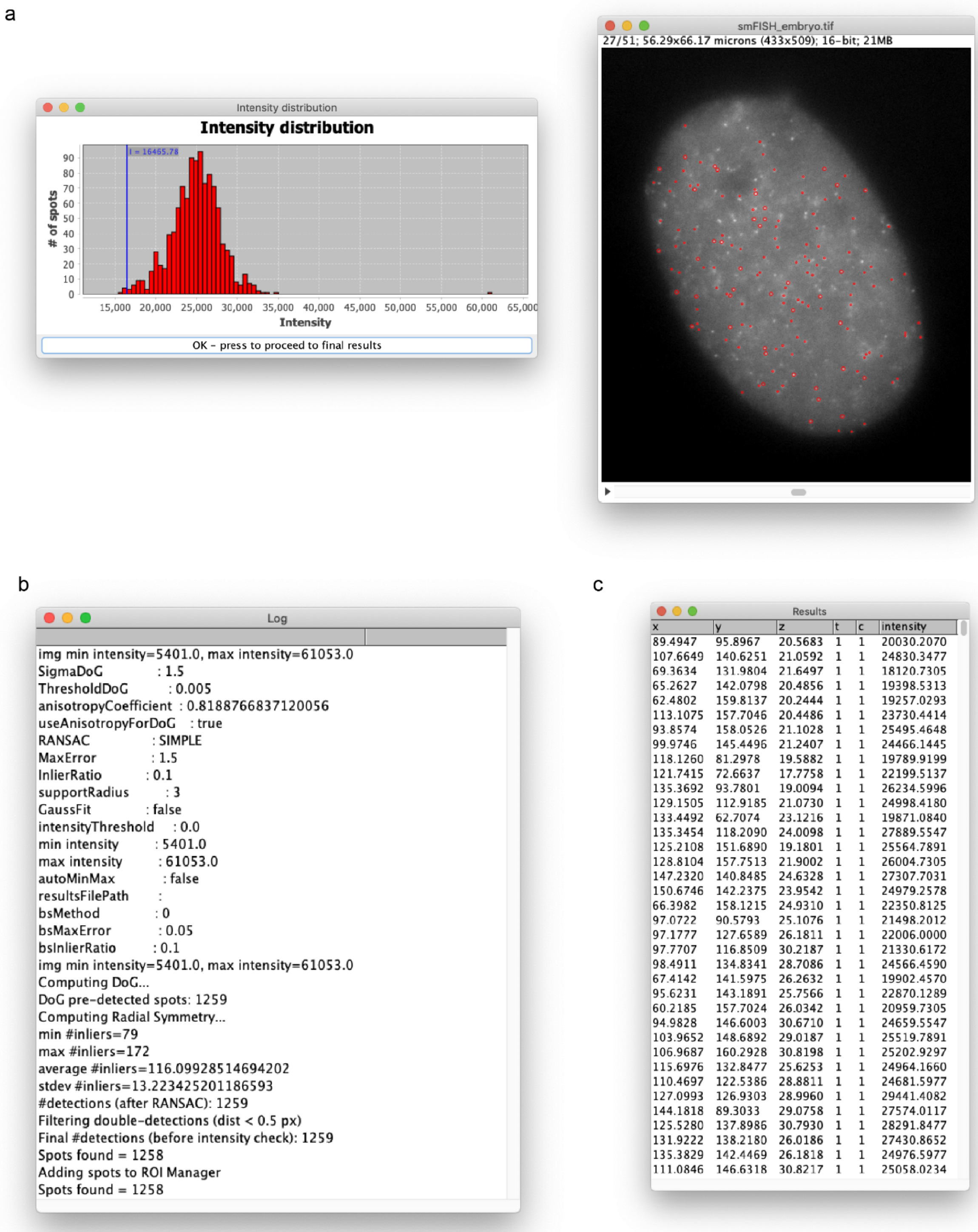


| x | y | z | t | c | intensity |
|---|---|---|---|---|---|
| 89.4947 | 95.8967 | 20.5683 | 1 | 1 | 20030.2070 |
| 107.6649 | 140.6251 | 21.0592 | 1 | 1 | 24830.3477 |
| 69.3634 | 131.9804 | 21.6497 | 1 | 1 | 18120.7305 |
| 65.2627 | 142.0798 | 20.4856 | 1 | 1 | 19398.5313 |
| 62.4802 | 159.8137 | 20.2444 | 1 | 1 | 19257.0293 |
| 113.1075 | 157.7046 | 20.4486 | 1 | 1 | 23730.4414 |
| 93.8574 | 158.0526 | 21.1028 | 1 | 1 | 25495.4648 |
| 99.9746 | 145.4496 | 21.2407 | 1 | 1 | 24466.1445 |
| 118.1260 | 81.2978 | 19.5882 | 1 | 1 | 19789.9199 |
| 121.7415 | 72.6637 | 17.7758 | 1 | 1 | 22199.5137 |
| 135.3692 | 93.7801 | 19.0094 | 1 | 1 | 26234.5996 |
| 129.1505 | 112.9185 | 21.0730 | 1 | 1 | 24998.4180 |
| 133.4492 | 62.7074 | 23.1216 | 1 | 1 | 19871.0840 |
| 135.3454 | 118.2090 | 24.0098 | 1 | 1 | 27889.5547 |
| 125.2108 | 151.6890 | 19.1801 | 1 | 1 | 25564.7891 |
| 128.8104 | 157.7513 | 21.9002 | 1 | 1 | 26004.7305 |
| 147.2320 | 140.8485 | 24.6328 | 1 | 1 | 27307.7031 |
| 150.6746 | 142.2375 | 23.9542 | 1 | 1 | 24979.2578 |
| 66.3982 | 158.1215 | 24.9310 | 1 | 1 | 22350.8125 |
| 97.0722 | 90.5793 | 25.1076 | 1 | 1 | 21498.2012 |
| 97.1777 | 127.6589 | 26.1811 | 1 | 1 | 22006.0000 |
| 97.7707 | 116.8509 | 30.2187 | 1 | 1 | 21330.6172 |
| 98.4911 | 134.8341 | 28.7086 | 1 | 1 | 24566.4590 |
| 67.4142 | 141.5975 | 26.2632 | 1 | 1 | 19902.4570 |
| 95.6231 | 143.1891 | 25.7566 | 1 | 1 | 22870.1289 |
| 60.2185 | 157.7024 | 26.0342 | 1 | 1 | 20959.7305 |
| 94.9828 | 146.6003 | 30.6710 | 1 | 1 | 24659.5547 |
| 103.9652 | 148.6892 | 29.0187 | 1 | 1 | 25519.7891 |
| 106.9687 | 160.2928 | 30.8198 | 1 | 1 | 25202.9297 |
| 115.6976 | 132.8477 | 25.6253 | 1 | 1 | 24964.1660 |
| 110.4697 | 122.5386 | 28.8811 | 1 | 1 | 24681.5977 |
| 127.0993 | 126.9303 | 28.9960 | 1 | 1 | 29441.4082 |
| 144.1818 | 89.3033 | 29.0758 | 1 | 1 | 27574.0117 |
| 125.5280 | 137.8986 | 30.7930 | 1 | 1 | 28291.8477 |
| 131.9222 | 138.2180 | 26.0186 | 1 | 1 | 27430.8652 |
| 135.3829 | 142.4469 | 26.1818 | 1 | 1 | 24976.5977 |
| 111.0846 | 146.6318 | 30.8217 | 1 | 1 | 25058.0234 |

**Supplementary Figure 4:** Thresholding the detected spots before the final results table. a) Option of thresholding the detected spots based on their intensity by changing the blue Intensity (I) cut-off. Spots that are currently within the selected threshold are marked with red circles in the z-plane with the highest intensity. b) The Log window with selected parameters and the total number of spots found. c) The Results table for all detected spots with their x,y,z location, and intensity.

### Advanced mode

In the **Advanced mode**, you can skip the interactive setting of parameters and only use already known parameters for the computation. After choosing **Advanced** in the first window (**Supplementary Figure 5a**), you will reach the second window to set the spot detection parameters (**Supplementary Figure 5b**). If you previously used the interactive mode to find the best parameters, they will be saved and set as default for the advanced mode. After you press **OK,** the computation is automatically done in all RS-FISH steps, and the same **Results table** as above is either saved or displayed.

### Scripting / headless

You can simply record the parameters you used for running RS-FISH (Plugins > Macro > Record). You can then apply it to a set of images using the Fiji/ImageJ macro language.

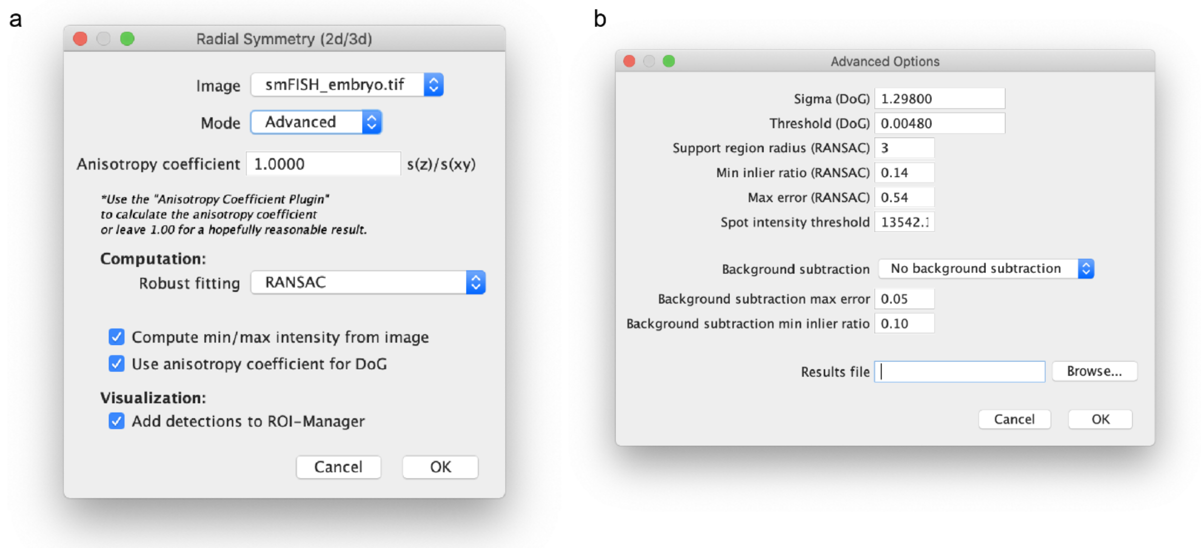


**Supplementary Figure 5:** RS-FISH advanced mode**.** a) To run the advanced mode of RS-FISH, select "Advanced" as running mode in the drop-down menu. b) The advanced options window allows you to select all detection parameters without manually adjusting them.

### Showing Detections

After RS-FISH computes all spots, the results table can be saved as CSV or directly be used to visualize the detected spots. There are three ways to visualize the RS-FISH detected spots:

- ROI Manager
- Fiji/ImageJ overlay
- BigDataViewer

If the option to transfer spots directly to the **ROI Manager** was chosen initially, the ROI manager would pop up with the results table in the last step (**Supplementary Figure 6a**).

The **Show Detections (ImageJ/Fiji)** plugin (Plugins > RS-FISH > Tools > Show Detections (ImageJ/Fiji)) can be used to overlay all spots stored in a CSV onto the current image for visual inspection of the final result using Fiji (**Supplementary Figure 6b**). The detected spots will be highlighted by red circles that are the largest in the z position where the center of the spot is.

a

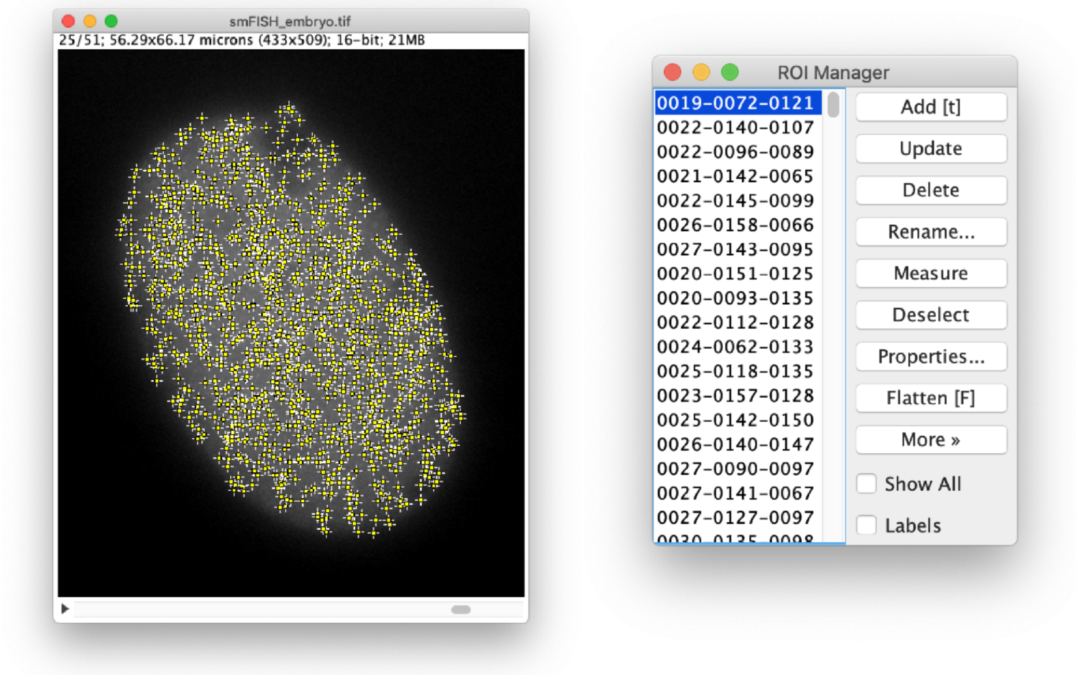


b

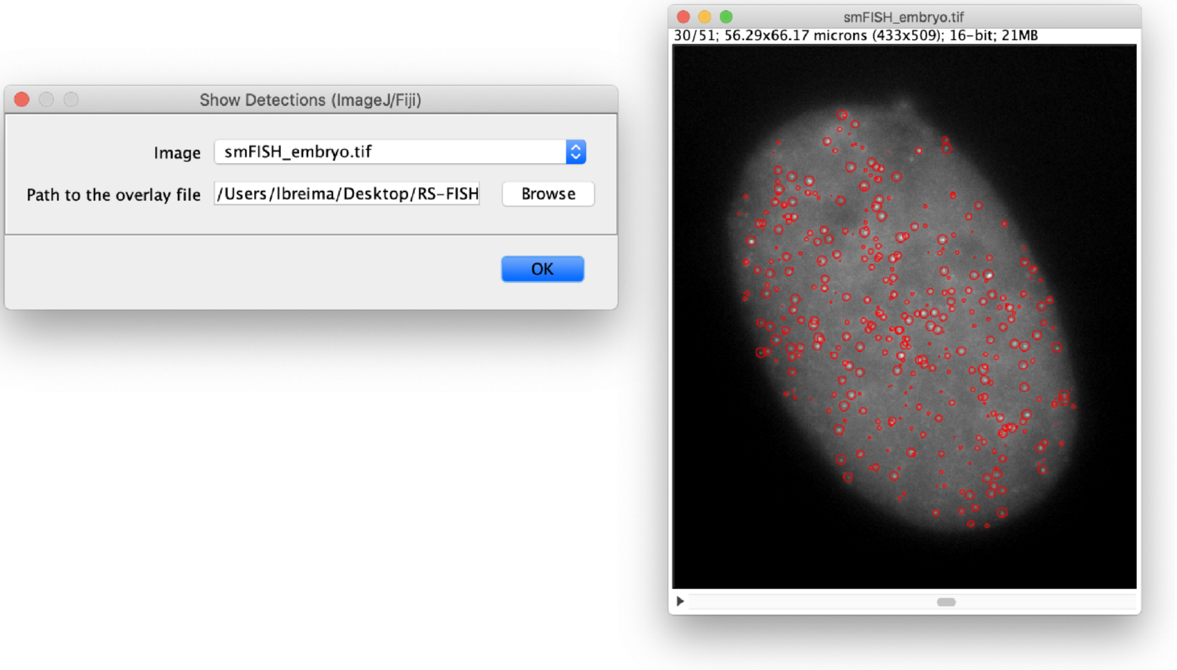


**Supplementary Figure 6:** Visualization options for detected spots. a) The detections can be transferred to the ROI manager for visualization or selection of single spots. b) Show detections using the RS-FISH plugin for a previously processed image.

The **Show Detections (BigDataViewer)** plugin (Plugins > RS-FISH > Tools > Show Detections (BigDataViewer)) can be used to visualize the spots using the BigDataViewer[8] (**Supplementary Figure 7**). In the first window are four options whether to open a saved image or CSV or use the currently active image and table (**Supplementary Figure 7a**). The intensity of the overlay points can be changed with the sigma slider.

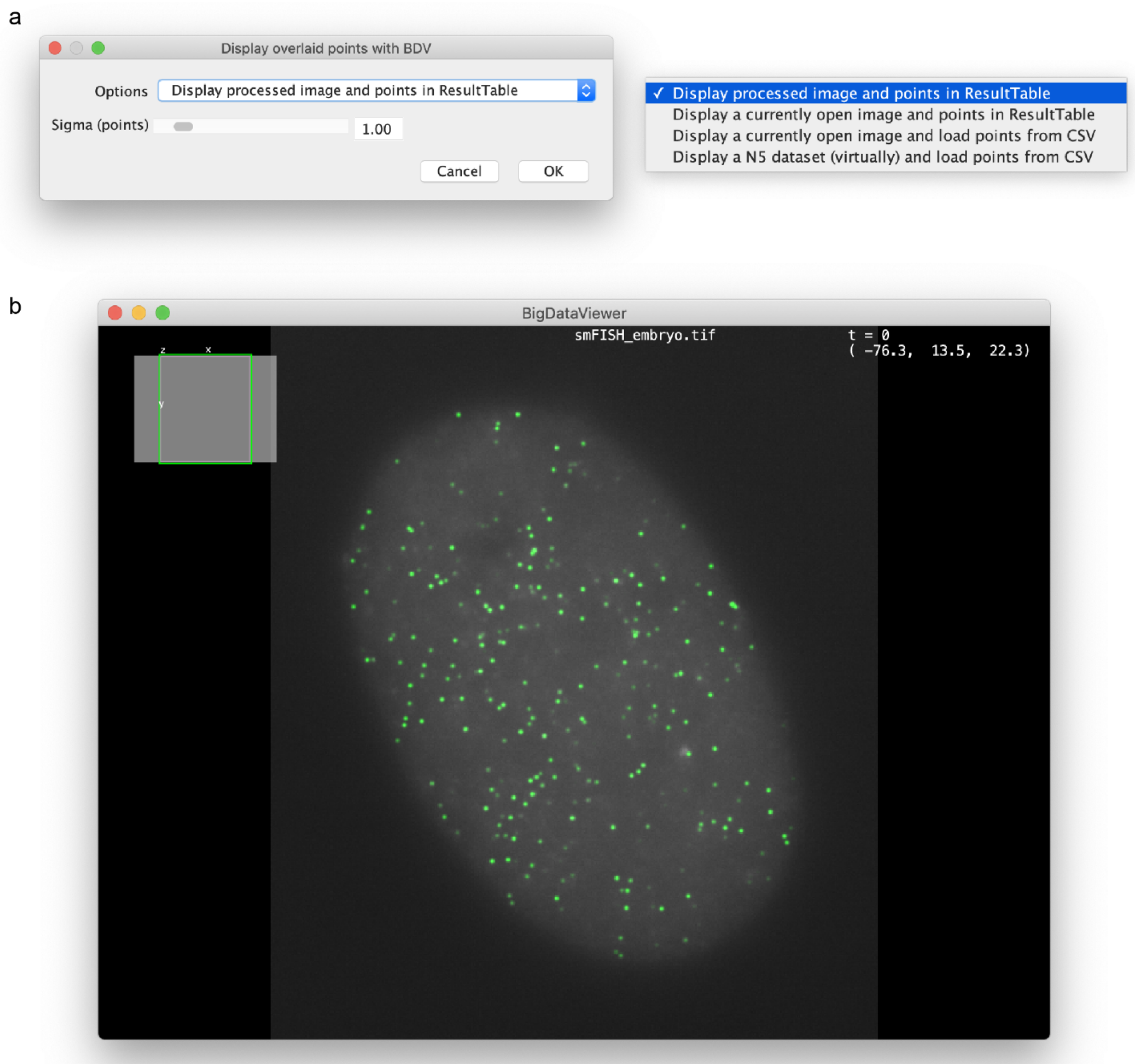


**Supplementary Figure 7:** Show detections using the BigDataViewer**.** a) Four options are available to display previously detected spots from either the open ResultsTable or a saved CSV. b) The BigDataViewer[8] displays detections (green) and the selected image (grayscale).

### Batch Processing and Macro Tutorial

RS-FISH plugin detections can also run from an ImageJ macro script from Fiji or headless from the terminal or a computing cluster.

This batch processing tutorial, as well as the example ImageJ macro script, can be found here:

https://github.com/PreibischLab/RS-FISH/tree/master/documents/example_scripts

A recommended procedure for running the plugin on a complete dataset is first running the plugin in Fiji in interactive mode on one (or a few) example images to find the best spot detection parameters. Then, use the parameters in a macro script that can be run from Fiji or headless.

Steps:
Find the best parameters for your dataset:

1. Open an example image from the dataset in Fiji.
2. Go to: Plugins > RS-FISH > RS-FISH
3. In the menu that opened - choose "Interactive" Mode, set your image anisotropy coefficient (z resolution / XY resolution), leave the rest as default values, then click ok.
4. Change the slider values of the parameters until you are happy with the detections you see on the image. Then click “Done”.
5. You can also change the intensity threshold in the “Intensity distribution” window according to the detections seen on the image. Once the correct threshold has been set, click “OK - press to proceed to final results”.
6. Save the Fiji log, as it details the parameters used. For this, in the “Log” window, go to: File > Save As, and save the Log.txt file at your chosen location.

Run in batch mode:

1. Open the Log.txt file you just saved, so you can copy the parameters you found in interactive mode to the macro script.
2. Open the `RS_macro.ijm` (in Fiji or a text editor) from the GitHub link above.
3. In the macro file, change the parameters (e.g., `anisotropyCoefficient`, `sigmaDoG`) at the beginning of the macro file (under the line `Define RS parameter`) to the values from the Log.txt file. Unless you are sure otherwise, only change the values of existing variables in the macro file.
4. In the macro file, it is recommended to keep the `useMultithread` variable as “use_multithreading” for a speedier run. This option is not available when RS-FISH is run in “Interactive” mode. If multithreading is used, `numThreads` should be set according to the number of threads on your machine. `blockSizX`, `blockSizY`, and `blockSizZ` should be set for chunking each image in the analysis to blocks. Note, different multithreading runs may result in ever so slightly inconsistent results.
5. In the macro file, change the `path` variable value to the parent directory of your .tif images (all tifs in all subdirectories will be processed).
6. In the macro file, change the `timeFile` variable value to the path you wish to save the running times file.
7. Call the script. It can be done from the Fiji GUI, from the terminal, or from a computing cluster. Example Linux command to run the macro script:

`<fiji_dir_path>/ImageJ-linux64 --headless --run </path/to/this/script>/RS_macro.ijm &> </path/to/where/you/want/yourlogfile>.log`

The macro above runs the same command as the command that is recorded when you record a run of the RS-FISH plugin in advanced mode.

Command Template:
`run("RS-FISH", "image=" + imName + " mode=Advanced anisotropy=" + anisotropyCoefficient + " robust_fitting=[" + ransacStr + "] use_anisotropy" +  " image_min=" + imMin + " image_max=" + imMax + " sigma=" + sigmaDoG + " threshold=" + thresholdDoG + " support=" + supportRadius + " min_inlier_ratio=" + inlierRatio + " max_error=" + maxError + " spot_intensity_threshold=" + intensityThreshold +  " background=[" + bsMethodStr + "] background_subtraction_max_error=" + bsMaxError + " background_subtraction_min_inlier_ratio=" + bsInlierRatio + " results_file=[" + results_csv_path + "]" +  " " + useMultithreadStr + " num_threads=" + numThreads + " block_size_x=" + blockSizX + " block_size_y=" + blockSizY + " block_size_z=" + blockSizZ;`

Example command:
`run("RS-FISH", "image=im.tif mode=Advanced anisotropy=0.6500 robust_fitting=[RANSAC] use_anisotropy  image_min=0 image_max=65535 sigma=1.203 threshold=0.0025 support=3 min_inlier_ratio=0.30 max_error=1.12237 spot_intensity_threshold=0  background=[No background subtraction] background_subtraction_max_error=0.05 background_subtraction_min_inlier_ratio=0.10 results_file=[/home/bob/Desktop/im_spots.csv] [use_multithreading] num_threads=40 block_size_x=128 block_size_y=128 block_size_z=16");`

Notably, running the tool/macro with a combination of parameters where sigma<1.5, threshold<0.002, and support>=3 will cause longer running times and require bigger memory, especially for bigger images.

### Distributed processing using RS-FISH-Spark

Distributed processing enables the analysis of terabyte-sized data or thousands of images. The Spark version of RS-FISH can analyze large N5 volumes in a distributed fashion locally, on the cluster, or in the cloud.

A tutorial on how to run a Spark-based version of RS-FISH can be found at https://github.com/PreibischLab/RS-FISH-Spark

### smFISH protocol for C. elegans embryos and larvae

This protocol for *C. elegans* embryos was adapted from the Raj lab [11] protocol with some modifications.

**Probe design**

All used smFISH probe sets (Custom Stellaris® RNA FISH Probe Set) were designed and manufactured by Biosearch Technologies using the Stellaris probe designer. Probes were designed to target exons and labeled with either Quasar 670, CAL Fluor Red 610, or Quasar 570. Typically far-red dyes performed better due to the autofluorescence of *C. elegans* embryos and especially in older stages.

**Preparation of worms for embryo collection**

Worms were synchronized by either bleaching or egg-laying and grown to the adult stage. At this step, it is important to have non-starved, healthy adult worms in sufficient amounts. Typically at least 5 x 6 cm plates full of worms were used.

**Collection of embryos**

For embryo collection, worms were washed off plates using M9 and collected in a 35 $\mu$m nylon filter. Worms were washed at least three times with H2O, then carefully transferred to a falcon tube using M9 and allowed to settle down. The supernatant was removed, and 5 ml freshly prepared bleaching solution was added to the worms. The dissolving of the adult worms was closely observed, and after 3 minutes, the tubes were spun at 3000 g for 1 min to collect the embryos. The supernatant was quickly removed, and the embryo pellet was vortexed. Then 10 ml of 1 x PBS-Triton were added, and the tube was centrifuged again for 3 min at 3000 g. This step was then repeated two more times until a clean embryo extract was left in the tube.

**Collection of larvae**

For the collection of L1 larvae, worms were synchronized by either bleaching or egg-laying and grown to the adult stage. Embryos were collected by bleaching as described above, and embryos were shaken in M9 overnight, and L1s were fixed the next morning.

**Fixing and permeabilization**

Embryos were resuspended in 1 ml fixation solution and incubated at RT for 15 min while rotating. Next, the tube was submerged into liquid nitrogen for 1 minute to freeze crack the embryo eggshells. The tube was then transferred to a beaker with RT water to thaw, and once it was fully thawed, it was kept on ice for an additional 20 minutes. After this incubation, the tubes were spun down at 3000 g for 3 minutes, and the supernatant was removed, and the embryos were washed twice with 1 ml of 1x PBS-Triton. The embryos were resuspended in 70% EtOH and kept at 4 °C for at least 24 hours. Embryos can be kept at 4 °C for at least several months.

**smFISH staining**

For smFISH staining of the previously fixed embryos, tubes were centrifuged at 3000 g for 3 minutes, and ethanol was carefully removed. The pellet can be pretty loose at this step, so removing the supernatant can also be done in two stages. Embryos were then resuspended in 1 ml wash buffer and vortexed. Tubes were centrifuged, as above, and the supernatant was removed. The embryos were then resuspended in a 50 $\mu$l hybridization solution, and 1 $\mu$l of each probe set (12.5 $\mu$M stock solution) was added directly to the sample. Tubes were then vortexed lightly and incubated at 37°C in the dark overnight. The next day, 0.5 ml of the wash buffer was added, the tubes vortexed and centrifuged to remove the supernatant. Next, 1 ml of wash buffer was added, and samples were incubated at 37°C for 30 minutes. After that, tubes were again centrifuged to remove the supernatant, and the embryo pellet was vortexed before adding 1 ml wash buffer. In this step, DAPI (5 ng/mL) was added to the wash buffer, and tubes were incubated at 37 °C for 30 minutes. The wash buffer was removed after centrifugation, and samples were washed once with 2x SSC.

**Mounting**

To mount the stained embryos, most of the liquid was removed from the tubes. About 15 $\mu$l of dense embryos (in 2x SSC) were used per glass slide and spread onto a coverslip (#1.5, 22 x 22 mm). The sample was left to dry for about 15 minutes, and then 15 $\mu$l of ProLong™ Diamond Antifade Mountant (Thermo Fisher) was added to the sample. The glass slide (SuperFrost, VWR) was pressed with the embryos and mounting media onto the coverslip. Slides were left at RT in the dark 24 hours before sealing the sides with nail polish and then stored at 4°C. Images were then acquired within two weeks of preparing the sample.

**Imaging**

Embryos were imaged on a Nikon Ti inverted fluorescence microscope with an EMCCD camera (ANDOR, DU iXON Ultra 888), Lumen 200 Fluorescence Illumination Systems (Prior Scientific), and a 100x plan apo objective (NA 1.4) using appropriate filter sets. Images were acquired with 90 z-stacks positions with 200 nm step-width using Nikon Elements software.

**Buffer**

- Collection buffer M9: 5.8 g $Na_2HPO_4$, 3.0 g $KH_2PO_4$, 0.5 g NaCl, 1.0 g $NH_4Cl$, Nuclease-free water to a final volume of 1000 ml
- 1x PBS (DEPC treated + autoclave + 0,05% of Triton X-100)
- Fixing buffer: 4% paraformaldehyde (PFA) in 1xPBS (DEPC treated + autoclave + 0,05% of Triton X-100)
- Bleaching stock solution: 2.5 ml 4N NaOH, 2.5 ml 5% NaClO, 5 ml Nuclease-free water, freshly made
- Washing buffer: 40 ml nuclease free water, 5 ml deionized formamide, 5 ml 20x SSC
- Hybridisation solution: 50 $\mu$l H20 (RNAse free), 37.5 $\mu$l EC 5 mg/ml, 25 $\mu$l formamide (at RT), 12.5 $\mu$l SDS (dissolved), 125 $\mu$l dextran sulphate 10 %
- 2x SSC

### smFISH protocol for mouse ES cells using cytospin

All pipetting steps were carried out at room temperature, and extra care was taken to never let cells dry in between buffer changes. Cells were grown in a 10 cm dish to 70-80% confluency and harvested by adding 1 ml warm trypsin. After 10 min, cells were carefully resuspended by adding 1 ml of warm ESC medium (DMEM (high glucose, without sodium pyruvate, cat no 41965062, Gibco, Life Technologies Ltd, UK ), supplemented with 15% FCS, 100 µg/ml penicillin/streptomycin, 2 mM L-Glutamine (Gibco, UK), 1x Non-essential amino acids (MEM-NEAA) (Gibco, UK), 50 µM beta-mercaptoethanol (Gibco, UK) and 10 ng/ml leukemia inhibitory factor (LIF, prepared in house)) and pipetting up and down several times. Cells were collected in a falcon tube with 4 ml ESC-medium and centrifuged at 1000 rpm on a Heraeus Multifuge L3 for 3 min (room temperature).

Cells were resuspended in 5 ml warm PBS, counted using a Neubauer chamber, and 1 ml of 4-6x$10^5$ cells per ml were transferred to a new falcon tube. To fix the cells, an equal volume of fixing solution (5 ml 10x PBS, 5 ml 37% formaldehyde, 40 ml H2O, in sample 2% PFA final concentration) and incubated for exactly 10 min at RT. Per slide, 100 µl of the solution was pipetted in an assembled cytofunnel (Shandon Single Cytofunnel, Thermo Fisher Scientific) and centrifuged in a Cytospin 4 (Shandon Cytospin, Thermo Fisher Scientific) at 1800 rpm for 3 min (with high acceleration setting). Cytofunnels were removed quickly, and slides were washed in 1x PBS for 5 min at room temperature.

To permeabilize the cells, slides were immersed in 70% ethanol and incubated at 4°C for 1 hour. Slides were washed in a washing buffer (5 ml 20x SSC, 5 ml formamide, 40 ml H2O) for 5 min at room temperature. During this step, the humidified chamber was assembled. A plastic container was lined with wet paper towels, and a slide holder was placed on top of it. To 50 µl room temperature, warm hybridization solution (100 mg/ml dextran sulfate and 10% formamide in 2X SSC) 0.5 µl of probe stock solution (12.5 µM) were added, mixed by vortexing and collected by a short centrifugation step. This resulted in a final probe concentration of 125 nM. A 50 µl drop of the probe containing hybridization buffer was added to the surface of a coverslip of the area where the cells were fixed. Slides were placed in the humidified chamber and incubated in the dark for 4 hours at 37°C.

Gently, slides were transferred to a fresh beaker containing a wash buffer after removing the coverslip. This washing step continued for 30 min in the dark at 37°C. Afterwards, the washing buffer was replaced by a DAPI-containing washing buffer (5 ng/ml) and incubated again in the dark at 37 °C for 30 minutes. The DAPI staining buffer was collected, and the final buffer, 2 X SSC, was added and incubated at room temperature for 5 minutes. Slides were mounted with 25 µl of Vectashield Mounting Medium, which was pipetted using a cut 200µl pipet tip. Excess fluid from the perimeter of the coverglass was gently wiped away, and the cover glass was sealed by clear nail polish to preserve the slides for a longer time. After 10 minutes of drying, slides were stored in a light-tight box at 4°C until being used for imaging.

Slides were imaged on a wide-field fluorescence microscope (DeltaVision Elite) using a 63x/1.4 N.A. plan-apochromat oil-immersion objective with N=1.514 oil (Applied Precision). The slides were illuminated with an SSI - LED light at different intensities. An Evolve EMCCD camera was used for image acquisition.

### smFISH protocol for cleared whole-mount adult Drosophila brains

A detailed description of the FISH method and FISH probes design were described previously (Long et al., 2017 [12]). Briefly, 3-5-day-old adult *Drosophila* were dissected in PBS and fixed for 55 min at RT in 2% paraformaldehyde. The tissues were then dehydrated and stored in 100% ethanol overnight at 4°C. After exposure to 5% acetic acid at 4°C for 5 min, the tissues were fixed again. The tissues were then incubated in PBS with 1% of NaBH4 at 4°C for 30 min followed by a 2h incubation in prehybridization buffer at 50°C (15% formamide, 2 × SSC and 0.1% Triton X-100). The tissues were transferred to 50 µL of hybridization buffer (10% formamide, 2 × SSC, 5 × Denhardt's solution, 1 mg/ml yeast tRNA, 100 µg/ml salmon sperm DNA, 0.1% SDS) with FISH probes against the targeted transcripts (50–100 ng/µL per reaction) and incubated at 50°C for 10 h and then at 37°C for an additional 10 h. After a series of wash steps, the tissues were dehydrated and cleared by xylene. The cleared tissues were mounted on a 1.5x3mm coverslip attached to the end of a 30 mm glass rod and imaged in a mixture of 90% 1,2-dichlorobenzene and 10% 1,2,4-trichlorobenzene medium.
